# A transcriptional patient map of systemic lupus erythematosus reveals disease-related multicellular immune programs conserved between blood and kidney

**DOI:** 10.64898/2026.04.28.721379

**Authors:** Jose Liñares-Blanco, Philipp Sven Lars Schäfer, Leoni Zimmermann, Ricardo Melo Ferreira, Daniel Toro Dominguez, Pedro Carmona Saez, Jovan Tanevski, Marta E. Alarcon Riquelme, Michael T. Eadon, Ricardo O. Ramirez Flores, Julio Saez-Rodriguez

**Affiliations:** European Molecular Biology Laboratory, European Bioinformatics Institute (EMBL-EBI), Hinxton, Cambridgeshire, U.K; Machine Learning lab in Life Sciences. Dept. of Computer Science and Information Technologies, Universidade da Coruña (CITIC), A Coruña, Spain; Heidelberg University, Faculty of Medicine, and Heidelberg University Hospital, Institute for Computational Biomedicine, Heidelberg, Germany; Department of Medicine, Indiana University School of Medicine, Indianapolis, IN 46202, USA; Unit of Inflammatory Diseases, Department of Environmental Medicine, Karolinska Institute, Nobel väg 13, 171 67, Solna, Sweden; Bioinformatics and Health Data Science. Pfizer-University of Granada-Andalusian Regional Government Centre for Genomics and Oncological Research (GENYO). Granada, Spain; Department of Statistics and Operations Research. University of Granada. Granada, Spain; Genetics and Genomics of Immune-Mediated Diseases, Centro Pfizer – Universidad de Granada – Junta de Andalucía de Genómica e Investigación Oncológica (GENYO), Granada, Spain

**Keywords:** Systemic Lupus Erythematosus, single-cell transcriptomics, multicellular coordination

## Abstract

Systemic lupus erythematosus (SLE) shows marked clinical and molecular heterogeneity, yet patient stratification often relies on gene expression signatures lacking multicellular context. Here we construct a transcriptional patient map of SLE by analyzing 1,167 total samples (783 SLE, 384 healthy controls) across different resolutions, including single-cell and bulk blood as well as spatially resolved kidney tissue transcriptomes. Using an unsupervised approach we inferred patient-level transcriptomic immune programs from two independent single-cell RNA sequencing cohorts of peripheral blood mononuclear cells (PBMCs), capturing both differences between SLE and health as well as within-SLE heterogeneity. Specifically, we identified four conserved programs comprising two multicellular inflammatory programs driven by interferon and TNF/NFkB activity across immune cells, and two cell type-specific programs reflecting CD8 T cell cytotoxicity and a CD4 T cell naive-to-effector state. Functional analysis of these programs revealed a rewiring of both cell-to-cell interactions and task allocation across cell types during disease activation. In addition, mapping these programs onto an external longitudinal blood transcriptomic cohort predicted flare risk and identified candidate blood protein biomarkers detectable by proteomics. Finally, we showed that these blood programs were enriched in immune-infiltrated glomerular regions from kidney biopsies of individuals with lupus nephritis using spatially resolved transcriptomic data, thereby linking systemic immune programs to local tissue pathology.

**Graphical Abstract:** 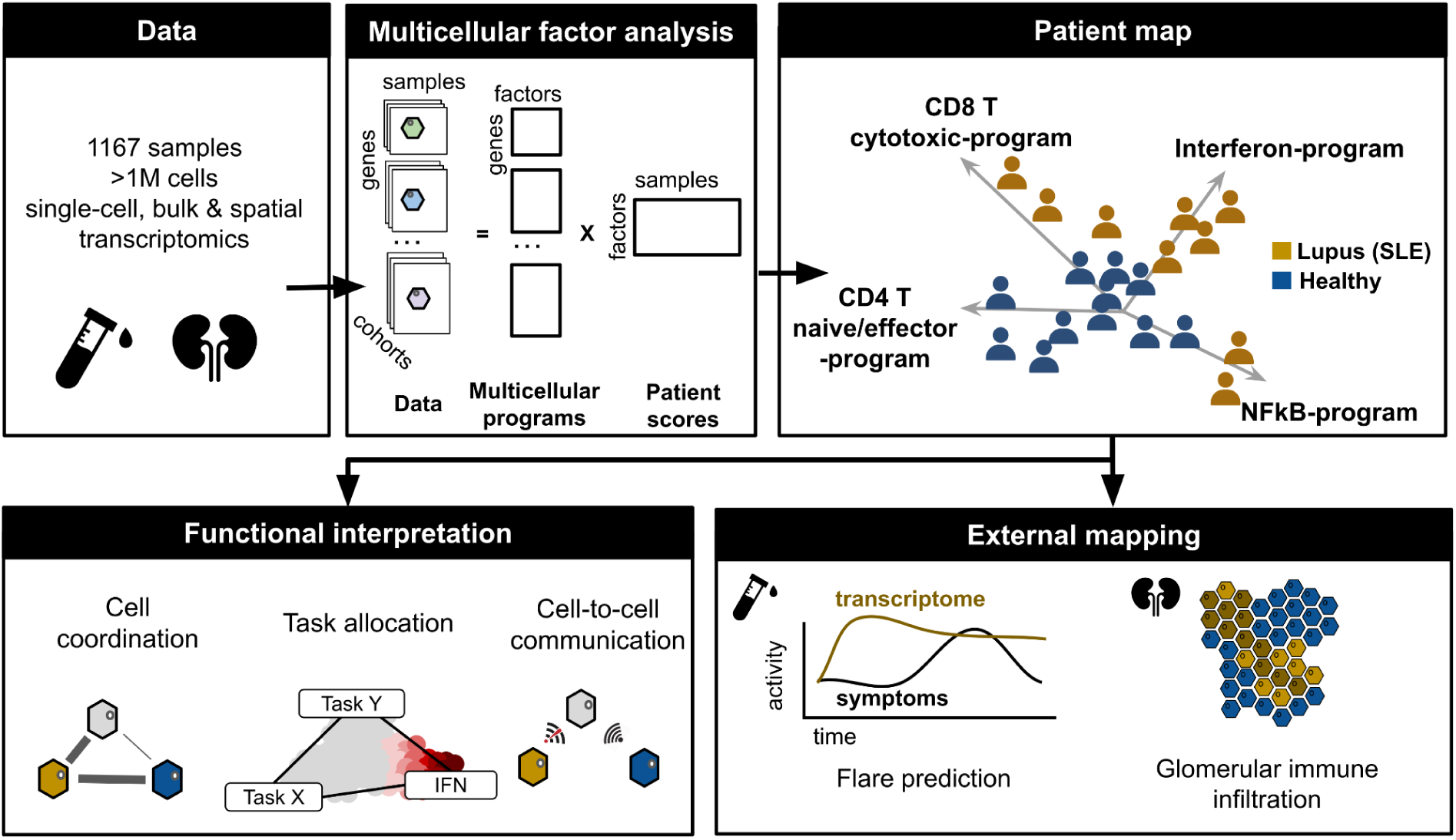

## 1. Introduction

Systemic lupus erythematosus (SLE) is an immune-mediated inflammatory disease characterized by pronounced molecular heterogeneity that only partly overlaps with clinical manifestations ^1,2^. Traditional clinical parameters and serological markers aid classification and monitoring but do not capture the diversity of immune activation states and pathogenic mechanisms driving individual patients^2–4^. Consistent with this complexity, multiple therapeutic strategies targeting both innate and adaptive immunity have been tested in SLE, yet only a limited number have shown consistent clinical benefit, and many patients fail to achieve durable responses or continue to flare under treatment^5^. Together, these challenges highlight the need to better characterize the heterogeneity of SLE.

Previous studies using bulk transcriptomic data from blood have shown that gene expression profiling can reveal molecular heterogeneity in SLE ^3,6–8^ . However, these approaches average signals across cell populations and therefore provide limited insight into cell state-specific variation. Single-cell RNA sequencing (scRNA-seq) of peripheral blood mononuclear cells (PBMCs) extends this exploration by enabling high-resolution profiling of gene expression at the level of individual cells. This approach has uncovered substantial cellular diversity ^9–13^, but prior analyses have largely focused on variation within predefined cell types rather than leveraging these data to more broadly characterize patient variability.

SLE as an autoimmune disorder can be studied through the lens of immune coordination, in which multicellular programs spread and sustain inflammation across cell types^14^. This motivates strategies that explicitly infer multicellular coordination from single-cell data ^15,16^, because analyzing each cell type in isolation can miss shared axes of variation that define patient-level immune states. Recent work^17^ has shown that coordinated multicellular programs can be inferred from single-cell data to stratify SLE patients and relate immune activity to clinical features. However, it remains unclear whether multicellular programs represent conserved, generalizable axes of disease biology that can be used to interpret existing bulk and spatial datasets and to connect systemic immune activation to organ-level pathology.

In this study, we analyzed 1,167 samples, including 783 SLE and 384 healthy controls, spanning three transcriptomic resolutions and two tissues. We leveraged PBMC single-cell RNA-seq profiles from two independent cohorts^9,10^ to infer patient-level multicellular programs that defined a transcriptomic patient map of SLE, enabling comparisons across individuals and characterization of inter-patient heterogeneity. In addition, we characterized cell-to-cell interactions emerging during active disease and the rewiring of transcriptional tasks within each cell type. We then mapped these programs onto external bulk cohorts ^3,18^, where they captured disease activity and predicted near-term flare risk. Finally, to link systemic immune coordination to organ pathology, we generated and analyzed spatial transcriptomes from lupus nephritis kidney biopsies and healthy kidney controls, and mapped blood-derived programs onto the tissue.

## 2. Results

### 2.1 Clinical, cellular and molecular comparison of single cell cohorts

To study immune cell coordination in SLE, we first curated two publicly available PBMCs scRNA-seq cohorts from Pérez et al. 2022 ^9^ and Nehar-Belaid et al. 2020 ^10^. The combined cohort comprised 316 samples, including 202 SLE cases and 114 healthy controls (**Sup. Fig. S1A**). First, we compared them at the clinical, immune cell composition, and transcriptional levels to define shared and cohort-specific variability.

Together, the cohorts span pediatric and adult patients (mean ages ∼16 years for pediatric and ∼42 years for adults; **Sup. Fig. S1B**) and showed the expected female predominance (93% female) (**Sup. Fig. S1C**). Among the 199 individuals with reported ancestry information, the main groups were Asian (n = 79), European (n = 59), Hispanic/Latino (n = 35), and African American (n = 19) (**Sup. Fig. S1D**). SLE Disease Activity Index (SLEDAI)^19^ was available for 184 SLE cases, ranging from remission to high activity (0–19) (**Sup. Fig. S1E**). Treatment exposure was heterogeneous: hydroxychloroquine was most common (n = 120), followed by oral steroids (n = 93), mycophenolate mofetil (n = 26), and methotrexate (n = 2) (**Sup. Fig. S1F**).

To enable comparison of immune cell composition and transcriptional profiles across studies, we harmonized cell type annotations between the two cohorts. We used the Pérez cohort as a reference and assigned cell-type labels to the Nehar cohort by mapping Nehar cells onto the Perez reference and transferring annotations based on similarity (see Methods). This procedure yielded a consistent set of 11 immune cell populations in both datasets: B cells (B), CD4⁺ T cells (T4), CD8⁺ T cells (T8), natural killer cells (NK), classical monocytes (cM), non-classical monocytes (ncM), conventional dendritic cells (cDC), plasmacytoid dendritic cells (pDC), plasmablasts (PB), proliferating T and NK cells (Prolif), and progenitors cells (Progen) (**Fig.1**A). To assess whether the cell type annotations were consistent across cohorts, we computed cell type–specific marker genes per cohort (**Sup. Fig. S2**A) and quantified cross-cohort concordance with the Jaccard index (**Sup. Fig. S2B**; Methods). Marker sets showed high, cell-type-specific concordance with Jaccard indices ranging from 0.37 to 0.74 (mean = 0.62), indicating that markers for a given annotation aligned with the same cell type and showed minimal overlap with other cell types.

**Figure 1:**
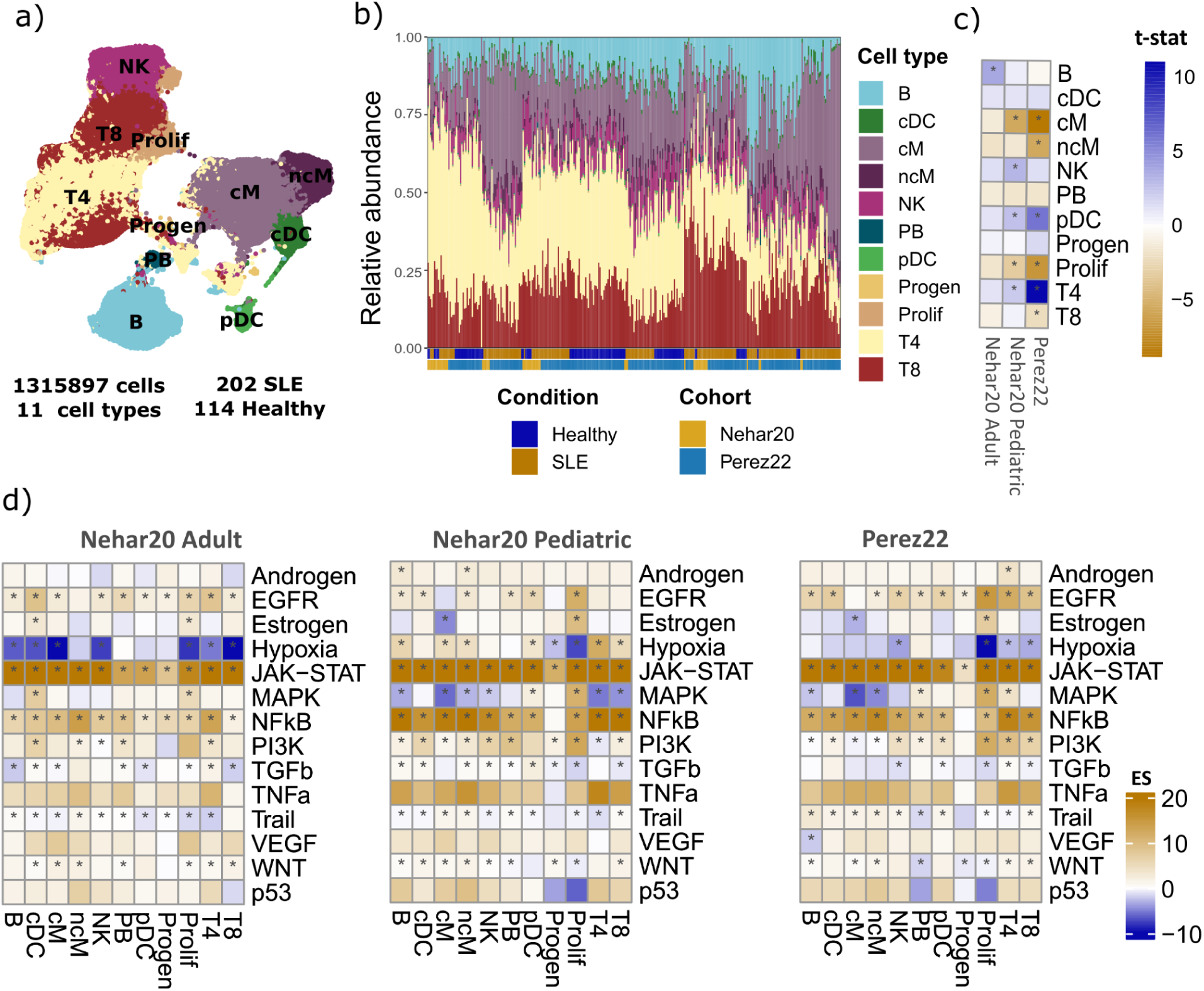
Single cell cohort comparison at clinical, cell type composition and molecular level. (A) Uniform Manifold Approximation and Projection (UMAP) of the integrated single-cell atlas (1,315,897cells); cells are colored by annotated cell type: B, CD4⁺ T (T4), CD8⁺ T (T8), NK, classical monocytes (cM), non-classical monocytes (ncM), conventional dendritic cells (cDC), plasmacytoid dendritic cells (pDC), plasmablasts (PB), T and NK proliferating cells (Prolif), and hematopoietic progenitors (Progen). (B) Hierarchical clustering of samples based on the composition of the eleven major immune cell types. (C) Differences in cell-type proportions between healthy and SLE across cohorts (two-sided *t*-test). Colors indicate effect direction (orange: SLE > healthy; blue: healthy > SLE). Asterisks denote significance (adjusted *P* < 0.1). (D) Pathway activity inference from DE *t*-statistics using PROGENy; rows, pathways; columns, cell types; values, signed activity scores (t-statistic from univariate linear model). Colors indicate enrichment direction (orange: SLE > healthy; blue: healthy > SLE). Asterisks denote significance (adjusted *P* < 0.05).

We calculated immune cell composition across samples and quantified the differences between healthy and SLE samples within each study. To account for potential differences between adult and pediatric SLE, the Nehar-Belaid et al. cohort was stratified into pediatric and adult subsets (**Fig.1B**). Cell-type proportion patterns were consistent across datasets (**Fig.1C**), with SLE samples showing higher proportion of classical and non-classical monocytes and T and NK proliferative cell types, and lower in CD4 T cells and plasmacytoid dendritic cell populations.

To assess molecular differences between SLE and healthy samples we calculated pseudobulk profiles by aggregating single-cell counts from all cells of the same type within each sample and performing differential gene expression within each cell type (see Methods). We defined significant genes (Wald test, adjusted *P* value < 0.05) across datasets and cell types and calculated the intersection (**Sup. Fig. S2C**). As expected by the larger sample size, the Pérez cohort yielded the highest number of differentially expressed genes across most cell types (average of 2,862 differential expressed genes per cell type), except for plasmablasts and progenitors, where the Nehar pediatric cohort showed a higher number (638 and 114 DEGs, respectively, compared to 272 and 15 in Perez).

Although the overlap of significant genes between cohorts was limited (mean pairwise Jaccarad Index = 0.11), the directionality of gene expression changes showed agreement. Pearson correlations of gene *t*-statistics across cohorts showed consistent trends (mean Pearson’s *r* = 0.34) (**Sup. Fig. S2D**), with the highest concordance for classical and non-classical monocytes (mean Pearson’s *r* = 0.47 and mean Pearson’s *r* = 0.49, respectively) and the lowest for progenitor cells (mean Pearson’s *r* = 0.085). To determine whether this agreement extended to higher-order signaling programs we inferred pathway activity in each cohort by evaluating the enrichment of PROGENy footprints^20^ using the differential expression *t-*statistics of each gene (**Fig.1**D; Methods). Inflammatory signaling pathways, particularly JAK–STAT and NF-κB, were consistently activated in SLE across datasets. Overall, both cohorts showed a conserved SLE-related transcriptional signal and motivated us to perform joint modeling to distinguish shared and cohort-specific immune activation patterns.

### 2.2 Immune multicellular programs define a transcriptional patient map of SLE

We applied multicellular factor analysis^16^ to identify coordinated transcriptional programs across immune cell types and to characterize sample-level immune heterogeneity in SLE. Briefly, the model decomposes variation between samples across multiple cell-types simultaneously by using their pseudobulk expression profiles. Analogous to principal component analysis, multicellular factor analysis decomposes the joint multicellular variability into a set of latent factors, each characterized by sample-level scores and gene loadings for every cell type. Gene loadings quantify the magnitude and direction of each gene’s contribution to a factor within a cell type, whereas factor scores summarize how strongly each sample expresses that factor. Because factors can capture both technical and biological sources of variation, we used the term multicellular programs to refer specifically to factors that are associated with known biological covariates.

Together, these factors define a shared low-dimensional space in which each sample can be positioned according to its multicellular immune profile, providing a transcriptional patient map of SLE heterogeneity.

We fitted the multicellular factor model on the combined PBMC scRNA-seq dataset. To reduce technical noise, we filtered the pseudobulk matrices to retain only well-represented cell-type–sample combinations and reliably detected genes, selecting highly variable genes per cell type while accounting for cohort-specific batches as the final input features (**Sup. Fig. S3A-B; Methods**). We restricted the analysis to nine immune cell types, excluding plasmablasts and progenitors because of their limited representation after quality-control filtering. We defined six groups corresponding to the processing batches reported in the original studies: the Nehar adult (N = 12) and pediatric (N = 44) subsets, and four processing batches in the Pérez cohort based on original publication (Pérez 1.0, N = 27; Pérez 2.0, N = 136; Pérez 3.0, N = 31; Pérez 4.0, N = 66). The composition of these groups with respect to disease status was unbalanced: Pérez 1.0 contains only healthy donors (N = 27), whereas the remaining groups include both SLE and healthy individuals in varying proportions (**Sup. Fig. S3C**). As a consequence, factors capturing disease-associated variation can only be estimated from groups that contain both healthy and SLE patients, while in healthy-only groups the same factors reflect baseline inter-individual variation.

The inferred model comprised 19 factors and captured, on average, 42.1% of the variance across cell types (CD8⁺ T cells highest, mean R^²^ = 60.1%; Prolif cells lowest, R^²^ = 26.6%; **Sup. Fig. S3D**). To characterize the main sources of variation captured by the factor space, we quantified its association with key clinical and demographic variables using multivariate linear regression. Disease condition and self-reported ancestry were the sources of variation better explained by the factors (R^²^ = 33.2% and R^²^ = 32.1%, respectively), followed by age (R^²^ = 18.8%) and sex (R^²^ = 7.4%) (**Sup. Fig. S3E**). As disease condition was the primary driver of factor variation, we focused subsequent analyses on factors linked to SLE biology.

To identify multicellular programs related to SLE variation we selected factors explaining >10% of variance in at least one cell type and one group. Five factors (1, 2, 3, 4, and 10) met this criterion (**Sup. Fig. S3F**). Factor 1 was broadly multicellular, explaining substantial variance across all nine cell types and in every group that contained SLE patients (up to 38.5% in classical monocytes), but was inactive in the healthy-only Pérez 1.0 group. Factor 2 explained variance in B cells, CD4⁺ T cells, dendritic cells, and classical monocytes, but predominantly within the Pérez 2.0 group, which contains the highest number of SLE samples. Factor 4 was cell-type-restricted, capturing variance exclusively in CD8⁺ T cells yet consistently across all six groups (22.0–34.4%). Factor 10 was similarly restricted, active primarily in CD4⁺ T cells in the two largest Pérez subsets (Pérez 2.0 and 3.0). Factor 3 was active almost exclusively in the healthy-only Pérez 1.0 group and was therefore excluded from subsequent analyses. Factors 1, 4, and 10 differed significantly between SLE and healthy individuals (two-sided t-test: P = 4.1 × 10^⁻¹³^, P = 6.7 × 10^⁻⁴^, and P = 2.4 × 10^⁻⁸^, respectively). Factors 1 and 4 were elevated in SLE, whereas higher Factor 10 scores marked a more healthy-like profile. In contrast, Factor 2, active predominantly in the Pérez 2.0 subcohort (N = 120 SLE, N = 16 healthy), captured within-disease heterogeneity rather than overall disease status (**Fig.2A, Sup. Fig. S3F**). We also confirmed that the remaining four factors preserved most of the association with SLE condition (*R²* = 23.6%) while decreasing associations with age (*R²* = 12.1%), self-reported ancestry (*R²* = 9.4%) and sex (*R²* = 2.0%) (**Sup. Fig. S3E**). These results suggest that our multicellular factor model is able to summarize patient variability in terms of four programs that capture both shared and cell-type-specific processes, and thus we decided to characterize their potential mechanistic and clinical implications.

**Figure 2:**
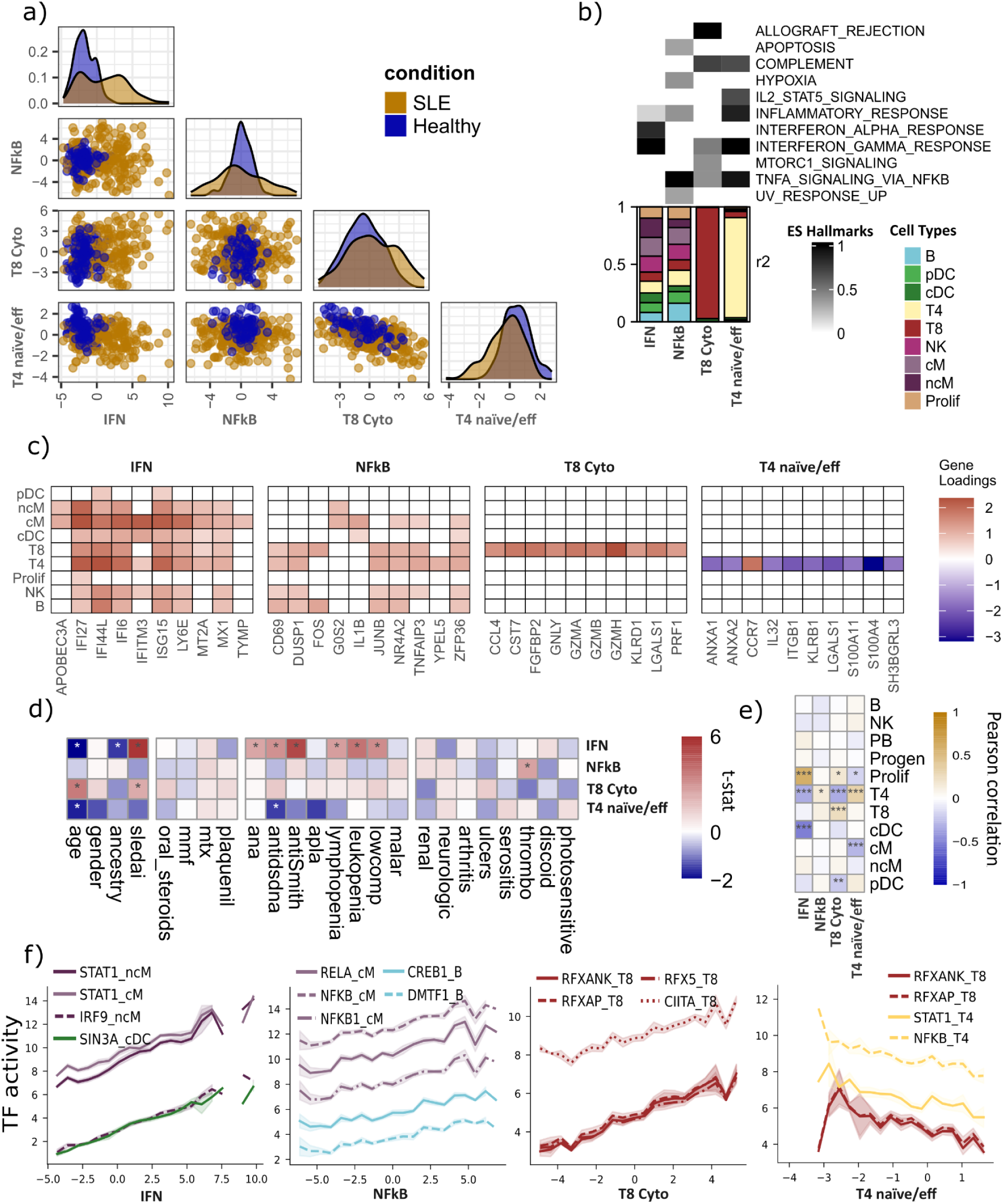
Multicellular programs in SLE. (A) Sample-level loadings for the four disease-relevant multicellular programs. Scatterplots show pairwise relationships between programs; diagonal panels show marginal density of program scores for SLE (orange) and healthy (blue) samples. (B) Coordination and functional annotation. Top: MSigDB Hallmark^35^ pathway enrichment for each program (absolute enrichment scores). Bottom: Explained variance per–cell-type for each program, highlighting multicellular versus cell-type–restricted programs. Explained variance is corrected to be relative to the total of each program. (C) Top contributing genes per factor across cell types. Heatmaps show gene loadings (top ten features by absolute weight). (D) Associations between factor loadings and demographic, clinical, and comorbidity variables in SLE. Tiles display *t*-statistics from linear models; asterisks denote significant associations (adjusted *P* < 0.1, ANOVA). (E) Relationship between program scores and immune-cell abundances across SLE individuals. Tiles show Pearson correlations with significance indicated by asterisks (adjusted *P <* 0.05). (F) Transcription factor activities associated with each program. Activities were inferred per sample and cell type using an univariate linear model with the CollecTRI regulon network^21^. For each program, the top transcription factors were selected based on the strongest linear association (slope) between program scores and TF activity across SLE samples. Lines show mean TF activity within bins along the program score axis; shaded areas represent the standard error of the mean. Colors indicate cell type.

### 2.3 Biological characterization of SLE-associated immune multicellular programs

To biologically characterize each SLE-associated multicellular program, we examined three complementary aspects of each program: explained variance, gene loadings, and sample scores. First, we quantified the variance explained in each cell type to distinguish multicellular programs, those explaining variance in at least two cell types, from cell type–specific programs, which primarily explain variance in a single cell type. Second, we used cell type–gene loadings to (i) define program marker genes based on the largest absolute loadings and (ii) perform pathway enrichment. Third, we treated program scores as per-sample measures of program activity and correlated them with sample-level features, including clinical data, immune cell composition, and inferred transcription factor activities using the CollecTRI^21^ gene regulatory network (**Fig.2B–F**; **Sup. Table 1**).

Program 1 displayed coordinated variability across all cell types (mean R² = 17.8%), with the strongest contributions from non-classical monocytes (26.8%), classical monocytes (26.1%), and conventional dendritic cells (22.1%) (**Fig.2B**). Pathway enrichment and top contributing genes indicated a type I interferon response, with interferon-stimulated genes (IFITM3, IFI27, ISG15, MX1) broadly expressed across myeloid and lymphoid lineages, which is in line with prior interferon signatures in SLE blood across cell types ^9,10^ (**Fig.2C**). It showed the strongest associations with clinical SLE features, including autoantibodies (ANA, anti-dsDNA, anti-Smith), cytopenias (lymphopenia, leukopenia), complement consumption (hypocomplementemia) and SLEDAI disease activity (**Fig.2**D). In addition, it correlated positively with the proportion of T and NK proliferating cells (0.9% of total cells; Pearson’s *r* = 0.60, P < 2.2 × 10⁻¹⁶) and negatively with CD4⁺ T cells (26.2% of total cells; Pearson’s *r* = −0.34, P = 9.2 × 10⁻⁷) and conventional dendritic cells (1.3% of total cells; Pearson’s *r* = −0.50, P = 2 × 10⁻¹⁴), but did not reflect correlation with cM or ncM proportions, pointing to interferon-driven reprogramming within existing monocytes rather than compositional shifts (**Fig.2E**). Consistently, transcription factor activity inference highlighted canonical interferon regulators (STAT1 and IRF9) mainly in monocytes (**Fig.2**F). These observations defined this program as a multicellular interferon program (IFN program) coordinated across all immune cell types and captures the dominant clinically associated molecular variation in SLE.

Program 2 showed moderate variability concentrated in B cells, classical monocytes, CD4⁺ T cells, and conventional dendritic cells, predominantly within the Pérez 2.0 group (R² = 10.6–13.9%; mean R² across all groups and cell types = 3.8%) (**Fig.2**B). Pathway enrichment pointed to TNF/NF-κB–mediated inflammation, and top genes such as IL1B and TNFAIP3 were characteristic of TNF/NF-κB–driven inflammatory responses ^22–24^, whereas the co-expression of early activation markers (CD69 and FOS) suggested an early lymphocyte activation program ^25,26^ (**Fig.2C**). No strong associations with clinical data (**Fig.2**D) nor with immune cell proportions (**Fig.2**E) were found for this program. Transcription factor analysis showed positive correlations with inflammatory regulators NFKB1 and RELA in classical monocytes and with CREB1 in B cells (**Fig.2**F). We interpreted this program as a NF-κB multicellular inflammatory program (NF-κB program) that captures variation in inflammatory activation among SLE patients, independently of IFN-related variation.

Program 4 showed cell-type–restricted variability, with variance explained predominantly in CD8⁺ T cells (mean R² = 27.7%; **Fig.2**B). Pathway enrichment highlighted allograft-rejection signatures, and the top contributing genes (GZMA, GZMB, PRF1, KLRD1) defined a cytotoxic effector module, consistent with canonical cytotoxic T cell programs ^27,28^ (**Fig.2**C). Clinically, this program displayed weaker associations than the IFN program but was significantly related to SLEDAI, indicating a contribution of this cytotoxic program to disease activity (**Fig.2**D). At the compositional level, this program correlated positively with CD8⁺ T cells (21.1% of total cells; r = 0.25, P = 4.2 × 10⁻⁴) and negatively with CD4⁺ T cells (26.2% of total cells; r = −0.34, P = 8.7 × 10⁻⁷), consistent with a compositional expansion of cytotoxic CD8⁺ T cells in SLE ^9,29–31^ (**Fig.2**E). Program scores correlated positively with inferred activities of RFX5, RFXANK and CIITA (core regulators of MHC class II transcription) consistent with prior evidence of CIITA inducibility in SLE CD8⁺ T cells^32,33^ (**Fig.2**F). Together, these results defined this program as a CD8⁺ T cell–restricted cytotoxic program (T8 cytotoxic program) linked to disease activity in SLE.

Program 10 was specific to CD4⁺ T cells in terms of variance explained, indicating another cell-type–restricted program (mean R² = 8.6% in CD4⁺ T cells; <0.5% in all other cell types) (**Fig.2**B). Top positive-loading gene (CCR7) was consistent with naive CD4⁺ T cell states^34^, whereas the strongest negative loadings (S100A4, LGALS1, ITGB1, KLRB1) indicated effector-differentiated CD4⁺ T cells, together defining an axis from naive to effector differentiation (**Fig.2**C). Higher program 10 scores, which reflect a more naive-like profile, were associated with decreased SLEDAI scores and with younger age. (**Fig.2**D). Compositional analysis revealed positive correlations with CD4⁺ T cell abundance (26.2% of total cells; r = 0.33, P = 2.1 × 10⁻⁶) and negative correlations with classical monocytes (27.4% of total cells; r = −0.29, P = 3.1 × 10⁻⁵) (**Fig.2E**). Consistent with a less inflammatory state, transcription factor analysis showed negative correlations between this program and STAT1 and NF-κB family members in CD4⁺ T cells, as well as with RFXAP and RFXANK in CD8⁺ T cells (**Fig.2**F). Therefore, this program was defined as a CD4⁺ T cell-restricted differentiation program (T4 naive/effector program) in which higher scores mark a preserved naive compartment and lower scores reflect effector differentiation, with the naive-skewed profile associated with lower inflammatory signaling and disease activity.

To validate cross-cohort generalizability, we projected the four programs onto an independent bulk transcriptomic SLE cohort (PRECISESADS cohort^3^ (N SLE samples = 226; see Methods) and found significant positive enrichment in the four previously defined molecular subgroups: IFN program in the Interferon cluster, NF-κB and T8 cytotoxic program in the Inflammatory cluster, and T4 naive/eff program in the both Lymphoid and Healthy-like clusters (**Sup. Fig. S3G)**. This cross-platform concordance confirmed that the multicellular programs captured reproducible axes of immune variation in SLE across cohorts and technologies.

Collectively, the four programs fall into two complementary categories. The IFN and NF-κB programs represent coordinated inflammatory responses that operate across multiple immune cell types, capturing distinct axes of innate immune activation in SLE, one dominated by type I interferon signaling and strongly linked to clinical manifestations, the other reflecting TNF/NF-κB-driven inflammation independent of IFN and capturing within-disease heterogeneity. In contrast, the T8 cytotoxic and T4 naive/effector programs represent cell-type-restricted shifts in lymphocyte differentiation states: an expansion of cytotoxic CD8⁺ T cells associated with higher disease activity, and a loss of the naive CD4⁺ T cell compartment that tracks with both age and disease severity.

### 2.4 Interferon-driven multicellular coordination in active disease

Having identified multicellular programs associated with SLE, we next leveraged the IFN program, to infer cell-type dependency networks that define the most general potential coordination processes between immune cell-types occurring during disease activation. Because this program is broadly shared across cell types and is a known hallmark of disease status, it provides a natural framework to ask which cell types co-activate and depend on each other when the IFN response is engaged. To do this, we computed cell-type–specific IFN program activity scores for each sample and used cross-validated ridge regression to predict each cell type’s activity from the activities of all other cell types, separately in SLE and healthy samples. Predictive performance of the models summarized how tightly each cell type’s activation is coordinated with the rest, while the regression coefficients captured directed dependencies between cell types, defining a cell-type dependency network (**Sup. Fig. S4A**; see Methods).

This analysis yielded two cell-type dependency networks, one for SLE and one for healthy samples, in which most cell types were well predicted by the activity of the remaining cell types (SLE mean R² = 0.93; healthy mean R² = 0.85), with the exception of pDCs and proliferating T and NK cells (Prolif), which were well predicted only in the SLE network (R² = 0.81 and R² = 0.89, respectively) but not in healthy network (R² = 0.50 and R² = 0.27) (**Fig.3A**). This suggested that, under disease conditions, pDCs and proliferating lymphocytes become tightly embedded in a coordinated IFN circuit, consistent with the known role of pDCs as major producers of type I IFN ^36,37^ and with the IFN-driven promotion of T and NK cell expansion ^38,39^. To identify coordination changes associated with disease, we compared cell type coefficients between the two networks and classified directed cell type dependencies as SLE-gained when present only in the SLE network, or healthy-specific when the reverse was observed (**Sup. Fig. S4B**, see Methods).

**Figure 3:**
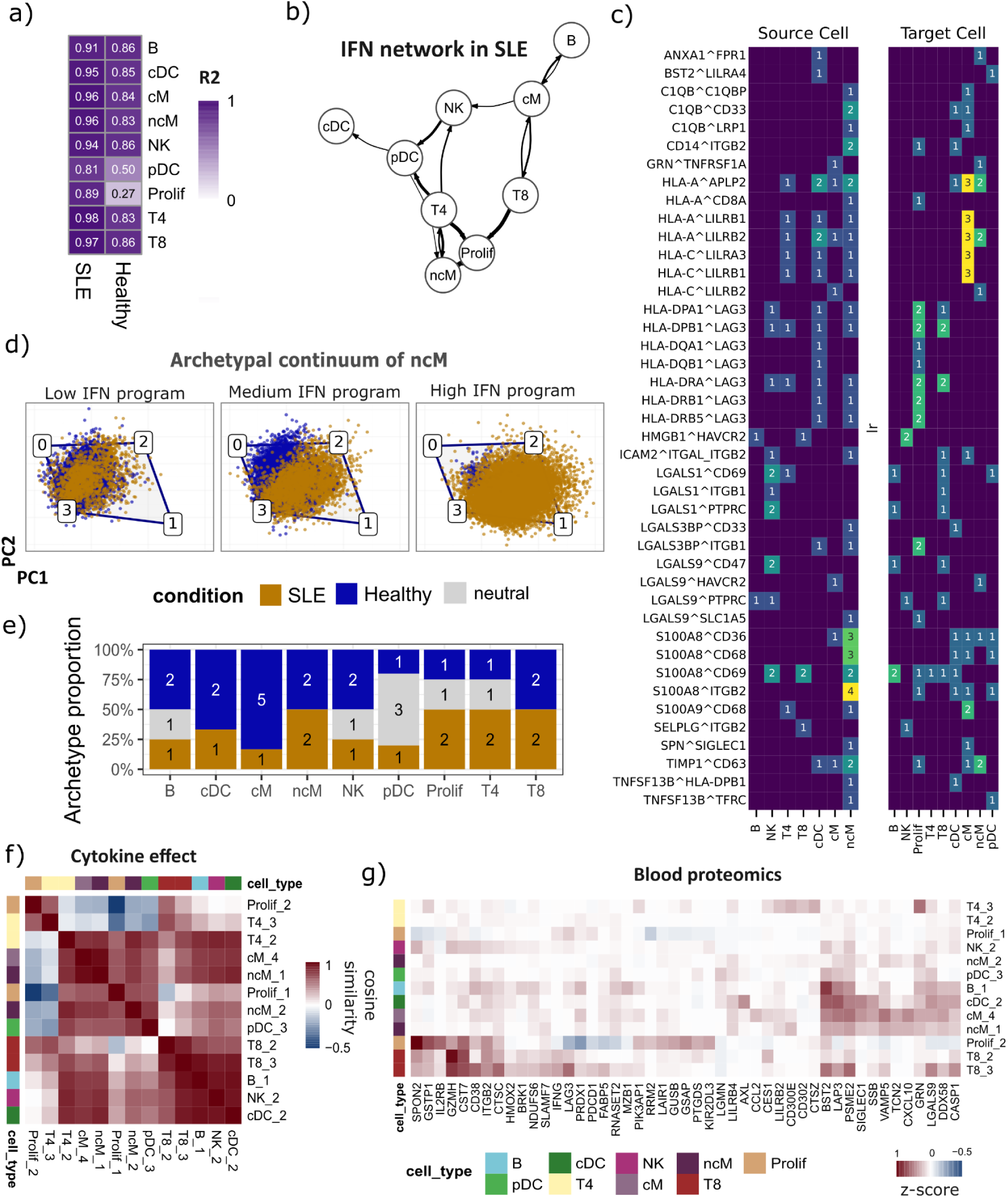
Interferon-driven coordination across immune cell types and its effect on task allocation. (A) Predictive performance (R2) to predict each of the immune cell types in both healthy and SLE networks. (B) SLE coordinated inferred network using IFN program. Edge thickness represents the predictive importance (betas from ridge regression model) between two cell types, while directionality indicates if the cell type was used as source or as target in the model. Darker and thicker connections denote stronger intercellular coordination. (C) Heatmap of all LR pairs significantly correlated with IFN program across cell types. Left and right panels indicate ligand-expressing (source) and receptor-expressing (target) cell types, respectively. Numeric values denote the number of significant interactions per combination (D) Archetypal analysis of ncM identifies four transcriptional archetypes (A0–A3). Scatter plots show ncM distributions across the archetypal space and along the IFN program by stratifying samples into terciles based on IFN program scores, and ncM from each tercile are visualized in the first two Harmony-corrected components. (E) Proportion of archetypes identified to be enriched along the IFN program per cell type. (F) Cosine similarity of SLE-associated archetypes based on cytokine-response footprint activities (see Methods). Rows and columns are hierarchically clustered (Ward’s D2 method). (G) Genes that are both upregulated in at least one SLE archetype (see Methods) and reported as elevated in lupus patient plasma in the Disease Blood Atlas. Rows represent SLE-associated archetypes and columns represent proteins detected in blood. Values correspond to archetype-level expression scores (z-scores). Rows and columns are hierarchically clustered (Ward’s D2 method).

Focusing on SLE-gained dependencies (**Fig.3B**), CD4⁺ T cells emerged as a central coordinator, with the strongest connections toward proliferating cells, pDCs, non-classical monocytes, and NK cells. Proliferating cells also displayed strong associations with non-classical monocytes. In addition, pDCs show SLE-specific outgoing associations toward conventional DCs and non-classical monocytes. Overall, these results suggest an SLE-specific coordination network centered on CD4⁺ T cells, proliferating lymphocytes, and non-classical monocytes.

These cell-type networks capture coordinated transcriptomic states, showing that when one cell type activates a program, others often do as well, but they do not identify the specific molecular interactions that could drive or reflect this coordination. We therefore investigated putative cell–cell communication (CCC) interactions associated with this IFN program. Using LIANA+ ^40^, we estimated, for each sample, CCC interactions across the nine cell types (see Methods). To focus on interactions most relevant to the observed coordination, we restricted the analysis to cell-type pairs connected in the dependency network and correlated each interaction score with IFN program activity across samples. This identified 94 significant interactions (Pearson’s r > 0.3, P < 0.05), corresponding to 42 unique ligand–receptor pairs involving 25 ligands and 26 receptors (**Sup. Fig. S4C, Sup. Table 2**).

To contextualize these IFN-associated interactions at the cellular level, we summarized, for each significant ligand–receptor pair, how often each immune cell type contributes the ligand (source) or the receptor (target) among 94 significant interactions (**Fig.3C**). First, non-classical monocytes emerged as the dominant sender population, contributing 40 of the 94 interactions, consistent with a myeloid-centered communication hub in IFN-high disease (**Sup. Fig. S4D**). The alarmin S100A8 was involved in 17 interactions and was most frequently sent by these monocytes, while one of the top-correlated events, SPN–SIGLEC1 (Pearson’s r = 0.55), further underscores IFN-driven myeloid activation, being SIGLEC1 (CD169) a canonical interferon-stimulated gene in monocytes. Second, we observed an antigen-presentation and checkpoint module centered on LAG3, being LAG3 the most frequently targeted receptor (17 of 94 interactions), predominantly paired with MHC class II ligands (HLA-DPA1, HLA-DPB1, HLA-DQA1, HLA-DQB1, HLA-DRA, HLA-DRB1, HLA-DRB5), with proliferating lymphocytes as the primary recipients followed by CD8⁺ T cells. Third, recurrent MHC class I–LILR interactions (HLA-A and HLA-C paired with LILRB1, LILRB2, and LILRA3) suggested a complementary inhibitory axis within the myeloid compartment, consistent with IFN-high conditions simultaneously engaging activation and counter-regulatory pathways. These results indicated that IFN-high SLE samples were characterized by a myeloid-centered communication architecture, anchored by non-classical monocytes as predominant senders, that coordinates alarmin and antigen-presentation signals with checkpoint and inhibitory receptor engagement on proliferating lymphocytes.

### 2.5 Archetypal analysis reveals rewiring of task allocation across cell types during disease activation

Disease activation in SLE involves coordinated transcriptional programs across multiple immune cell types, yet how these programs are partitioned into specialized functions within individual cells remains unclear. To address this, we used archetypal analysis to decompose single-cell gene expression profiles into a small number of extreme transcriptional states, referred to as archetypes (see Methods). We interpret these archetypes as candidate functional tasks, such that each cell’s position relative to these archetypes reflects its bias toward one or a combination of these tasks. We then asked how disease activation, quantified by the IFN program, redistributes cells across archetypes and thereby alters task allocation within each population^41–43^. For each archetype, we tested whether patients with higher IFN program activity harbor cells that are closer to that archetype by regressing patient-level archetype distance against IFN program scores (OLS, Benjamini–Hochberg FDR correction; see Methods). Archetypes were classified as SLE-associated (negative coefficient, FDR < 0.05), or healthy-associated (HC) (positive coefficient, FDR < 0.05), or neutral (**Sup. Fig. S5A**).

We first applied this approach to non-classical monocytes, previously identified having a central role in the SLE-associated network. Archetypal analysis resolved ncM into four archetypes **(Fig.3D)**. Two archetypes were classified as SLE-associated and two as HC-associated (**Fig.3D, Sup. Table 3**). The first SLE archetype (ncM_1) showed tight coupling with the IFN program (R² = 0.81) and was enriched in interferon-alpha (t = 51.8) and interferon-gamma response (t = 48.2), defining a canonical IFN-driven state (**Sup. Fig. S5B**). Explaining less variance (R² = 0.13), the second SLE archetype (ncM_2) was characterized by TNF-alpha/NF-κB signaling (t = 30.0), inflammatory response (t = 17.4), and IL-6/JAK/STAT3 (t = 9.8), defining a complementary inflammatory program that is partially decoupled from the IFN axis.

Extending this analysis across all nine cell types, we identified at least one SLE-associated archetype per population, yielding 13 disease-associated tasks in total, with the dominant SLE archetype consistently enriched for interferon pathways **(Sup. Fig. S5B**). However, the distribution of these archetypes varied across immune cell types (**Fig.3E)**. Some populations (B, NK, cDC, cM, pDC) exhibited a single SLE-associated archetype, whereas others (ncM, T4, T8, proliferating cells) resolved into two distinct disease-associated archetypes. We observed that disease activation in some cell-types induced a loss of multiple healthy-associated archetypes that transitioned to a single dominant SLE-associated archetype (cDC, cM), while in other cell-types we observed that some archetypes remained shared between healthy and SLE conditions on top of enriched disease archetypes (NK, B, pDC, Prolif, T4). This suggests that disease activation rewires transcriptional task allocation to different extents across immune cell types.

To understand which cytokine signals shape these disease-associated archetypes, we scored the activity of 90 cytokines in SLE-associated archetypes by using a single-cell cytokine dictionary of human peripheral blood^44^, a reference map of cytokine-driven transcriptional responses across peripheral immune cell populations (see Methods). Clustering cytokine-response profiles of SLE-associated archetypes showed that archetypes group by shared cytokine sensitivity rather than by cell type of origin. (**Fig.3F**). Major functional differences between SLE and HC archetypes based on enrichment analyses were associated with a pan-interferon module (IFN-α, IFN-β, IFN-γ, IFN-λ) and gamma-chain cytokines (IL-2, IL-7, IL-15) as the most strongly SLE-enriched signals, while HC archetypes were enriched for IL-10 and TGF-β1 signatures, suggesting that disease progression involves not only the gain of pro-inflammatory programs but also the loss of an immunoregulatory cytokine axis (**Sup. Fig. S5C, Sup. Table 4**). Together, these results showed that disease-associated archetypes enrich along a continuous gradient of IFN program activity, reflected cell-type-specific patterns of task redistribution and variability, but could be described by a shared cytokine response landscape across immune cell types.

### 2.6 Multicellular program activity predicts flare risk and identifies blood biomarker candidates

So far, we characterized these multicellular programs as biologically coherent axes of coordinated immune activity. For these programs to be clinically useful, two conditions must be met: their activity should be detectable through accessible measurements, and they should associate with meaningful disease outcomes.

To address the first, we asked whether the transcriptional states captured by SLE-associated archetypes leave a detectable trace in the peripheral blood proteome. For each archetype, we defined a gene signature based on its top upregulated genes (see Methods) and matched these against proteins significantly elevated in SLE patient plasma from the Disease Blood Atlas^45^, identifying 47 proteins with concordant transcriptional and proteomic evidence (**Fig.3G**). These proteins were distributed across transcriptionally distinct cell populations, suggesting that archetypal analysis can help trace the cellular origin of circulating biomarkers in SLE, and that program activity may have a clinically accessible proteomic readout.

To address the second, we mapped the programs onto an independent longitudinal bulk transcriptomic cohort of SLE patients (n = 352 visits from 84 patients; median 4 visits per patient) and asked whether they capture clinically relevant variation in disease activity. For each visit, we computed multicellular program activities by combining cell-type–specific activities, giving more weight to cell types that contributed more strongly to each program (see Methods). We confirmed the association of these programs with SLE Disease Activity Index (SLEDAI)^19^ using linear mixed-effect models (FDR < 0.05; **Sup. Fig. S6A**). The IFN program showed the strongest positive associations across all cell types, followed by the NFkB program in NK, ncM, and proliferating cell types. Consistent with patterns observed in the single-cell cohort, the T8 cytotoxic program was positively associated with SLEDAI, whereas the T4 naive/effector program showed an inverse association.

We then compared program activity between visits in clinical remission (SLEDAI < 3) and visits with active disease (SLEDAI ≥ 3) (**Sup. Fig. S6B**). Program activity was significantly higher during active disease across all cell types, with the exception of the T4 naive/effector program. However, we observed high variability in remission visits, raising the possibility that elevated program activity during clinical remission reflects a subclinical immunologic state that precedes disease reactivation. To test this, we stratified remission visits (SLEDAI < 3) into short- and long-term remission based on whether they were followed by a flare (defined as a change in SLEDAI ≥ 4) within three- and six-month intervals (**Fig.4A**, Methods).

**Figure 4:**
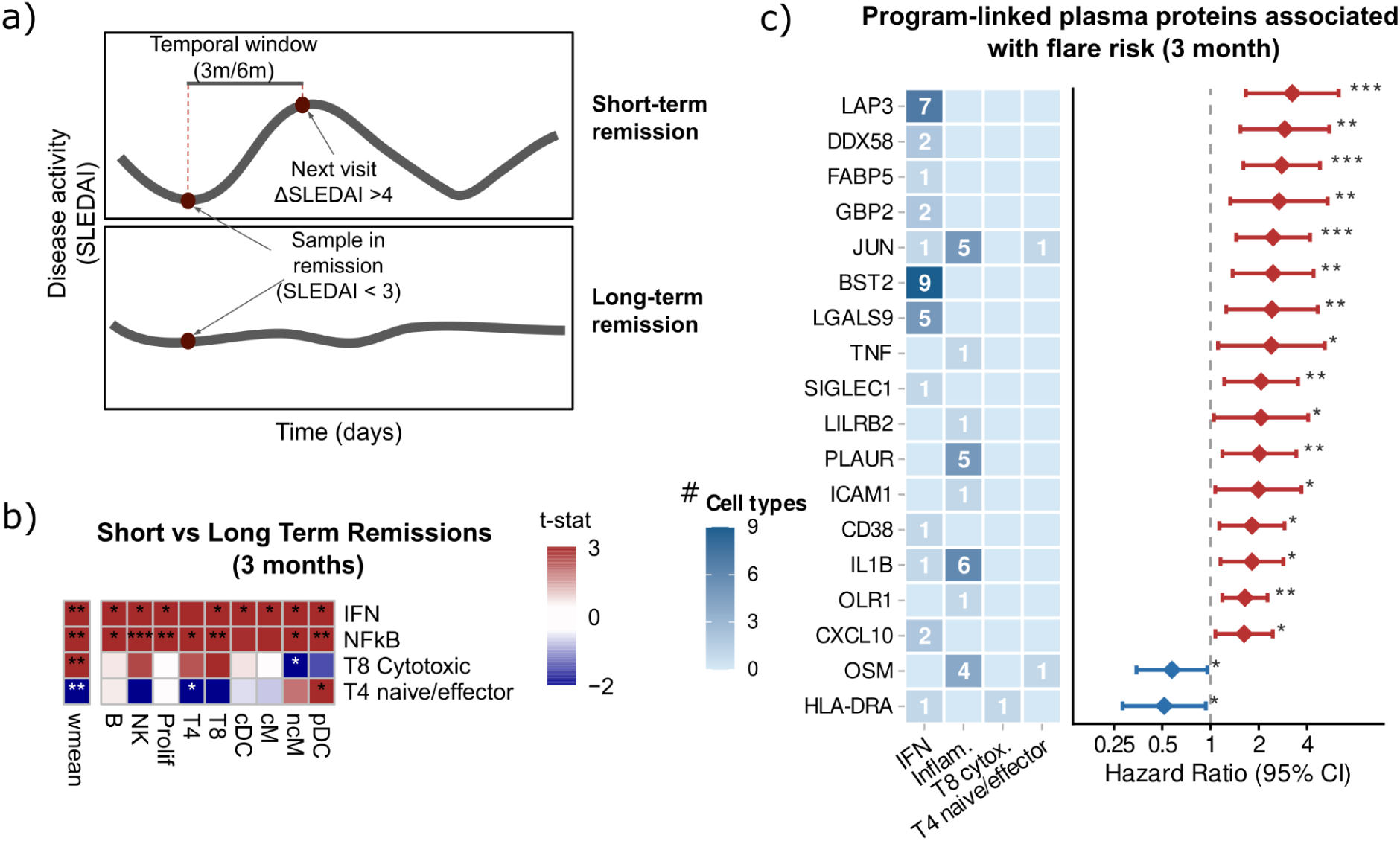
Multicellular programs anticipate flare in remission samples. (A) Schematic of the longitudinal labeling strategy used to stratify remission visits. Visits in clinical remission are defined by SLEDAI < 3 and are classified based on disease activity at the next follow-up within a predefined temporal window of 3 or 6 months. Remission visits followed by a flare at the next visit, defined as an increase in disease activity of ΔSLEDAI ≥ 4, are labeled as short-term remission, whereas remission visits not followed by flare within the window are labeled as long-term remission. (B) Heatmaps showing differential immune program activity between remission visits that preceded a clinical flare (short-term remission) and those with sustained disease control (long-term remission), at a 3-month prediction time frame. For each program–cell type combination, differences were assessed by two-sample t-test; colour represents the t-statistic, with positive values indicating higher program activity in short-term (pre-flare) visits. The left row ("wmean") summarises a weighted mean across cell types, where weights are proportional to the variance explained (R²) by each factor in each cell type as estimated by the multicellular factor model. P-values were corrected for multiple comparisons using the Holm procedure; asterisks denote significance: *p < 0.05, **p < 0.01, ***p < 0.001. (C) Combined plots showing genes significantly associated with flare risk at a 3-month prediction time frame, that overlap with top gene signatures of the multicellular programs and were detected in blood proteomic cohort upregulated in SLE patients. Left panel (heatmap): for each gene, the number of cell types in which it constitutes a top gene of the respective program (IFN, NFkB, T8 Cytotoxic, T4 naive/effector). Right panel (forest plot): hazard ratio (HR, 95% CI) from univariate Cox proportional hazards models, estimating the association between gene expression and time to flare. Proteins are ordered by HR. Red indicates proteins associated with increased flare risk (HR ≥ 1, p < 0.05); blue indicates protective associations (HR < 1, p < 0.05); grey indicates non-significant associations. Asterisks denote significance: *p < 0.05, **p < 0.01, ***p < 0.001.

Within the three-month window (16 short-remission visits, 59 long-remission visits), the IFN, NFkB and T8 cytotoxic programs were elevated in visits preceding a flare, whereas the T4 naive-effector program was higher in visits that did not precede a flare (adjusted P-value < 0.05, two-sample t-test; **Fig.4B**). Both IFN and NFkB programs showed consistent differences across multiple cell types, reinforcing the multicellular nature of this pre-flare state. Cox proportional hazards modelling further confirmed that NFkB was the dominant independent predictor at three months, capturing the signal of the remaining programs in pairwise models (**Sup. Table 5**). At six months, this pattern shifted: only the IFN program reached significance, predominantly at the multicellular level, while NFkB lost independent predictive value (**Sup. Fig. S6C, Sup. Table 5**). Together, these results suggest that proximal flare risk is driven by a broad multicellular inflammatory state involving both IFN and NFkB activity, whereas longer-term risk reflects a more diffuse systemic IFN tone, consistent with NFkB representing an acute inflammatory signal^46^ and IFN a chronic predisposition to disease reactivation^47^.

Finally, building on the proteomic overlap identified earlier, we extended this analysis to all four programs and asked whether individual genes driving each program carry predictive value for flare risk and are detectable in peripheral blood. Matching top-loading genes from each progam–cell-type against the Disease Blood Atlas (see Methods), we found several candidates significantly associated with flare risk in the longitudinal cohort, with genes linked to the NFkB program predominating at three months and IFN-associated candidates emerging as stronger predictors at six months(**Fig.4C**, **Sup. Fig. S6D**).

Together, these results showed that multicellular programs predicted flare risk during clinical remission, with NFkB activity dominating short-term prediction and IFN emerging as the stronger predictor at longer time frames, and identified individual blood-detectable genes linked to each program as candidate biomarkers

### 2.7 Spatial mapping of multicellular programs reveals immune-infiltrated glomerular niches in lupus nephritis

Multicellular programs identified in peripheral blood captured systemic immune activation. Therefore, they could inform on the presence of immune cell infiltration within affected organs. To test this, we focused on the kidney, a primary target in SLE where immune infiltration can lead to lupus nephritis. We generated and analysed an independent external cohort comprising three slides of 10x Visium spatial transcriptomics data from kidney biopsies of SLE patients with lupus nephritis and seven slides from healthy controls (**Sup. Fig. S7A**).

First, cell type abundance at each spot was estimated by deconvolution using a kidney single-cell reference^48^, and multicellular programs were mapped to then calculate spatial co-localisation between program activity and cell type abundance (**Fig.5A**, Methods). We observed how immune cells showed no significant spatial co-localisation with any program in healthy tissue, whereas in SLE biopsies they exhibited significant positive co-localization with IFN, NFkB and T8 cytotoxic programs (Moran I > 0.1, BH-adjusted p < 0.001; **Sup. Fig. S7B**). Specifically, B cells and myeloid dendritic cells displayed the strongest positive co-localization with the IFN and T8 cytotoxic programs (Moran’s I = 0.130 and 0.132, respectively; BH-adjusted p < 0.001), followed by non-classical monocytes (**Fig.5B**). B cells showed the largest disease-associated spatial shift, with co-localization nearly absent in healthy kidney (Moran’s I = 0.01) but increased in SLE (Moran’s I = 0.13) consistent with accumulation of B cells within inflammatory renal niches in lupus nephritis ^49,50^.

**Figure 5:**
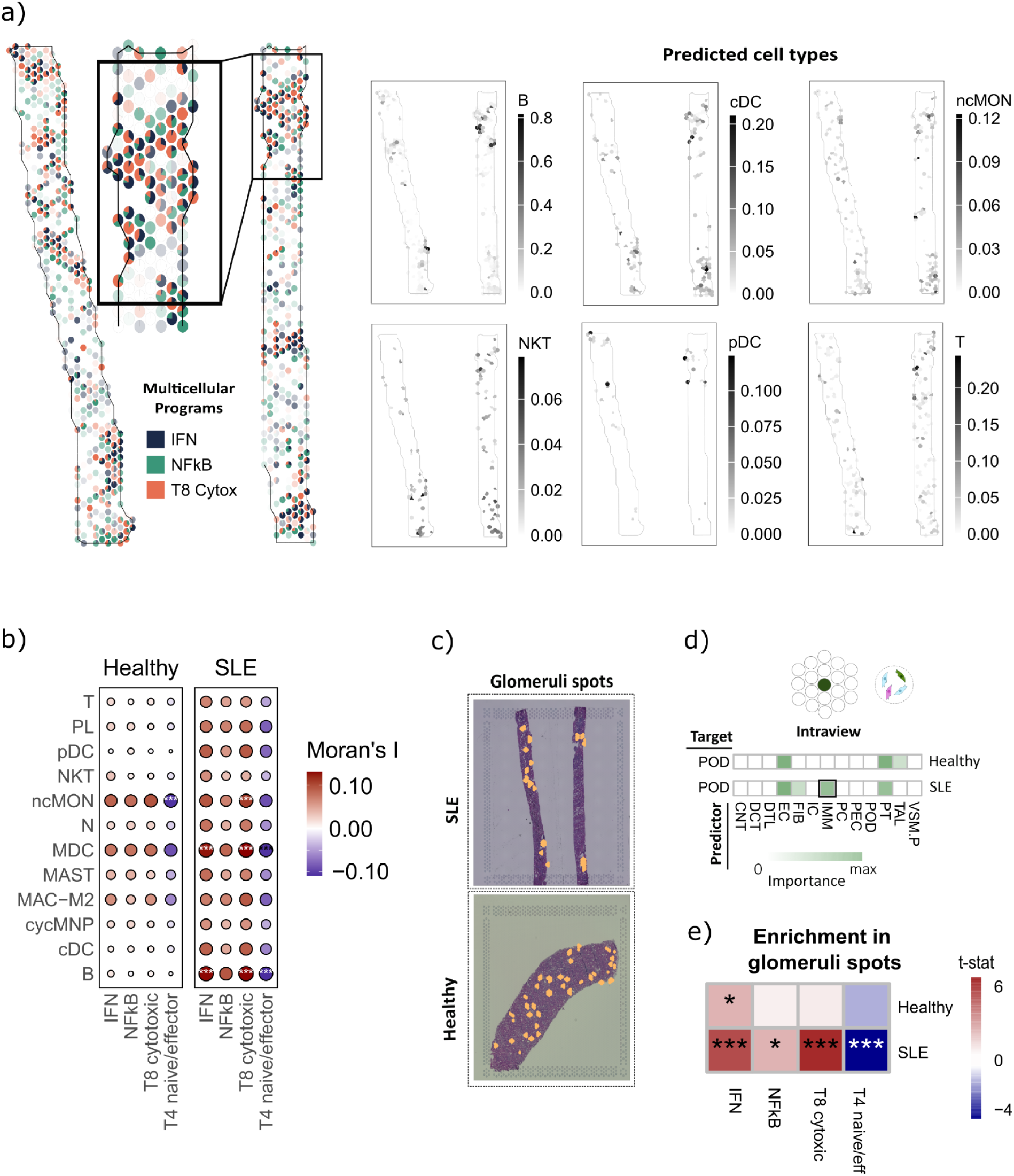
Spatial mapping of programs reveals immune cell infiltration in kidney. (A) Spatial mapping of selected multicellular programs (Factors 1, 2, and 4) onto kidney Visium sections. *Left*, representative SLE tissue section showing spatial distribution of program activity across spots. Each spot is represented as a pie chart displaying the relative contribution of the three factors, while color intensity indicates the mean factor score (opaque colors denote higher overall activity). *Right*, predicted spatial distribution of major immune cell types inferred using a Seurat-based deconvolution approach. Immune cell enrichment patterns highlight localized immune infiltration within the tissue. The single-cell reference used for spatial deconvolution was derived from the KPMP single-nucleus RNA-seq dataset^48^. (B) Spatial correlation between multicellular program activities and predicted immune cell-type abundances between conditions, quantified using bivariate Moran’s I. Moran’s I values were averaged across slides per condition (SLE, Healthy), and statistical significance was assessed by combining per-sample permutation-derived p-values using Fisher’s method, followed by Benjamini-Hochberg FDR correction. Dot size represents the magnitude of spatial association (|Moran’s I|); color indicates directionality (red = positive, blue = negative). Asterisks denote associations that are simultaneously statistically significant (FDR < 0.05) and exceed a minimum effect size (|Moran’s I| ≥ 0.10). (C) Example of glomeruli annotation and distribution across slides (SLE and healthy). (D) Feature importances from the intraview MISTy model predicting podocyte and composition within glomerular regions. The model captures spatial colocalization patterns by quantifying how strongly each cell type contributes to the prediction of others across spots. Results were aggregated separately for healthy and SLE slides. (E) Enrichment of programs across healthy and SLE glomerular spots. Tiles show *t*-statistics from linear models, with color indicating directionality of change (red = enriched in SLE, blue = enriched in healthy). Asterisks denote statistically significant differences (adjusted *P* < 0.05). Immune cell states annotations based on ^48^: B, B cells; PL, plasma cells; T, T cells; NKT, natural killer T cells; MAST, mast cells; MAC-M2, M2-like macrophages; cycMNP, cycling mononuclear phagocytes; MDC, myeloid dendritic cells; cDC, classical dendritic cells; pDC, plasmacytoid dendritic cells; ncMON, non-classical monocytes; N, neutrophils.

Given the spatial correlation of immune cell types with multicellular programs, we further inspected if immune cell infiltration occurred specifically within glomerular structures, where immune-mediated inflammation directly impacts renal filtration function in lupus nephritis. Thus, we focused on annotated glomerular regions (glomerular spots and neighbours) and applied a multi-view spatial modelling^51^ to predict the abundance of one cell type based on the abundance of the other cell types (see Methods). This framework allowed us to inspect contributions of cell types co-localized within the same-spot and from neighboring spots, enabling discrimination between direct co-localization and spatial proximity effects. Since podocytes are the major cell component of glomeruli, we predicted podocyte abundance, and observed that immune cell abundance was informative in SLE samples, but not in healthy samples, only when considering cell abundance within the same spot, (**Fig.5D**, **Sup. Table 6**), suggesting co-localization of immune and podocyte populations within glomerular regions in SLE, whereas contributions from neighboring regions were not detected.

Finally, we examined whether the multicellular programs were enriched in these immune-infiltrated glomerular niches. We restricted the analysis to annotated glomeruli spots and performed enrichment analyses comparing glomerular spots from SLE (n = 78) and healthy controls (n = 130). Again, we observed IFN (t-stat = 5.90), NF-κB/TNF (t-stat = 2.73) and T8 cytotoxic (t-stat = 7.07) programs enriched in SLE glomeruli, whereas the T4 naive/effector program was reduced (t-stat = -5.34) (**Fig.5E**). These results showed that the multicellular inflammatory programs identified in peripheral blood were recapitulated within glomerular niches in lupus nephritis kidneys, suggesting that the systemic immune activation captured in blood is also present within the glomerular compartment, where it may play a role in driving renal injury in lupus nephritis.

## 3. Discussion

In this study, we inferred multicellular transcriptional programs from single-cell RNA-seq data of PBMCs cells, and showed that sample-level immune variation captured by these programs was reproducible across independent bulk transcriptomic cohorts and detectable in spatially resolved kidney tissue from individuals with lupus nephritis. In total we combined data from 783 SLE and 384 healthy samples across multiple transcriptomic resolutions to study SLE variability. First, we applied unsupervised multicellular factor analysis to PBMC single-cell transcriptomes from SLE and healthy individuals and inferred four programs that captured coordinated transcriptional variation across immune cell types. These included two multicellular inflammatory programs driven by interferon and TNF/NF-κB activity, and two cell-type-restricted programs reflecting CD8 cytotoxicity and a CD4 naive-to-effector axis. Rather than assigning patients to discrete categories, these programs placed each individual along continuous, partially independent dimensions that could coexist within the same patient, forming a quantitative patient map. The continuous and quantitative nature of this map enabled the integration of other data modalities, and their correlation with clinical variation and disease activity, providing a system-level view of SLE immune activation.

Previous bulk and single-cell studies had defined molecular signatures of SLE and catalogued disease-associated immune states, and recent work had shown that multicellular programs could stratify patients and inform clinical outcomes^17^. In this study, we extended those results by learning programs in an integrated multi-cohort setting to separate shared from cohort-specific variation. This allowed us to identify new multicellular programs such as an NF-κB program that was partially decoupled from the IFN program and carried distinct prognostic information, for example, being an independent predictor of imminent flare risk. In addition, multicellular programs were coupled to within-cell-type archetypal analysis to map patient-level axes back to single-cell state variation. We further nominated blood-detectable gene markers linked to these multicellular programs and showed that the programs generalized across different data modalities, including blood proteomics, bulk transcriptomics, and spatially resolved kidney tissue, where they mapped onto immune-infiltrated glomerular regions, linking systemic immune activation to local tissue pathology. Together, these results positioned multicellular coordination as a complementary framework to cell state discovery that supported quantitative, transferable program-level stratification across datasets, modalities, and tissues.

Inference of cellular dependencies from multicellular programs provided a systems-level view of immune coordination in SLE and highlighted which cell types were most engaged in disease-associated activation. Focusing on the IFN program, the dominant axis of variation, we observed that interferon did not act only as a cell-intrinsic inflammatory signal in SLE, but as a multicellular organizing axis that coordinated state changes across immune compartments. This coordination appeared to be only partially explained by inferred cell–cell communication, indicating that systemic inflammatory coupling in SLE likely extended beyond canonical ligand–receptor signalling alone ^52^. At the same time, the interaction modules identified here pointed to candidate paired biomarkers that might capture coordinated sender–receiver states more precisely than single markers, thereby offering a potential route toward more accurate molecular monitoring. Importantly, this coordinated IFN response was associated with a cell-type-specific reallocation of cellular tasks rather than a uniform shift across all populations^42,53^, with some cell types showing pronounced state compression and others retaining or diversifying their functional repertoire. This heterogeneity may have important therapeutic implications, suggesting that treatments targeting systemic inflammatory drivers such as interferon may not uniformly reverse disease-associated task allocation across immune cell types. Cell-type-specific persistence of specialized inflammatory states may therefore contribute to variable treatments responses and underscores the need for more cell-type-aware immunomodulatory strategies.

A central translational challenge in SLE is that conventional clinical variables do not capture disease biology with sufficient resolution, which results in limited precision for patient stratification and therapeutic decision-making. This underscores the need for robust and reproducible molecular biomarkers that more precisely reflect the underlying immune processes ^54^. In this context, multicellular programs appeared to provide translational value beyond their original discovery setting, not only by generalizing across external cohorts but also by identifying states of subclinical immune activation that were not resolved by standard clinical indices ^55,56^. This raised the possibility that molecular remission may be more informative than clinical remission alone. The observation that different programs were associated with distinct temporal patterns of flare risk further suggested that inflammatory axes may operate across different prognostic time frames, with potential implications for time-sensitive monitoring strategies. Importantly, because key components of these programs were detectable in peripheral blood and were also linked to tissue-level pathology, these signatures provided a plausible bridge between systemic immune state and organ-level disease, as was previously reported^12^. Taken together, this supported the idea that multicellular programs could form the basis of practical biomarker frameworks for longitudinal monitoring, risk stratification, and earlier intervention in SLE.

This work has several limitations. First, our analysis captures only the transcriptomic component of SLE heterogeneity. Different processes, including genomic and epigenetic regulation, autoantibody specificity, and metabolic rewiring, may drive SLE pathogenesis, but can not be directly captured from transcriptomics. Other data modalities, including metabolomics and (epi)-genomics are likely to help us explain further patient-to-patient variability and may refine the programs described here. Since not all molecular variability is necessarily clinically actionable, a major translational challenge will be to identify which axes of variation are most informative for patient stratification, treatment selection, and longitudinal monitoring. Second, disease characterization across multi-cohort data remains vulnerable to partial confounding by cohort structure, batch effects, sample handling, unbalanced sampling, and clinical covariates such as medication exposure or intercurrent infection. Although harmonization and external projection mitigate some of these concerns in our patient map, given the data structure, we can not avoid including residual confounding information during inference. Third, coordination is inferred here from co-variation in gene expression across cell types and therefore does not establish causality or directionality. The SLE-gained dependencies and ligand–receptor modules should thus be viewed as hypothesis-generating candidates for experimental validation rather than definitive mechanistic circuits. Fourth, despite including both adult and pediatric samples, we did not identify robust age-specific factors, which may reflect limited statistical power or, alternatively, the predominance of shared inflammatory axes that obscure subtler developmental differences. Finally, both the size of the renal spatial cohort and the resolution of Visium spots limit generalizability and mechanistic inference. Spatial co-localization depends on deconvolution and reference atlases, does not resolve direct cell–cell contacts, and will require validation in larger cohorts and in higher-resolution spatial kidney datasets.

In conclusion, our patient map provides a complementary framework by representing each individual as a combination of coordinated immune programs rather than a single dominant signature. The patient map and program scores are publicly available, allowing external single-cell, bulk, and spatial datasets to be projected into a common coordinate system for cross-cohort comparison and integrated clinical and tissue-level interpretation. This will be particularly useful in the context of SLE management^57,58^, which increasingly combines standard immunosuppression with targeted therapies and steroid-sparing approaches^59,60^, but many patients still show variable responses and frequent flares^61^, highlighting the need for more robust patient stratification methods.

## 4. Methods

### Processing single cell transcriptomic studies

Single-cell RNA-sequencing count matrices were obtained from two publicly available studies: Pérez et al.^9^ (GSE174188) and Nehar-Belaid et al.10 (GSE135779). To harmonize cell type annotations across cohorts, we first used the Pérez dataset as a reference. Raw counts from Pérez were preprocessed following the original publication ^9^ (including gene/cell filtering, normalization, batch correction, and regression of technical covariates), and Scanpy’s^62^ *ingest* procedure was then applied to project Nehar-Belaid cells into the reference space and transfer cell-type labels using k-nearest-neighbor assignment to reference cells. This step yielded a shared cell type annotations across cohorts, including B cells, NK cells, plasmablasts (PB), progenitors (Progen), proliferating cells (Prolif), CD4⁺ T cells (T4), CD8⁺ T cells (T8), conventional dendritic cells (cDC), classical monocytes (cM), non-classical monocytes (ncM), and plasmacytoid dendritic cells (pDC).

Once cell type labels had been assigned, quality control filters were applied to retain genes expressed in at least three cells and cells with detectable expression of at least 200 genes. Clinical metadata were harmonized by keeping only covariates shared between cohorts. After this filtering, the Pérez et al. cohort comprised 957,596 cells across 25,275 genes from 261 samples, whereas the Nehar-Belaid et al. cohort comprised 358,301 cells across 24,435 genes from 56 samples. Throughout the manuscript, we refer to these quality-controlled, label- and metadata-harmonized count matrices as the preprocessed scRNA-seq datasets.

### Processing bulk studies

We analyzed three bulk transcriptomic datasets: the PRECISESADS RNA-seq cohort ^3^ and one longitudinal microarray cohort^18^ (GSE121239, Affymetrix HT HG-U133+ PM Array Plate)

#### PRECISESADS (RNA-seq)

Raw count expression values were processed in edgeR^63^. A DGEList object was constructed and lowly expressed genes were removed using filterByExpr (using diagnosis as the grouping variable). Library size and compositional differences across samples were corrected by trimmed mean of M-values (TMM) normalization (calcNormFactors, method = “TMM”). Normalized expression values were converted to log-counts per million (logCPM) using cpm(log = TRUE, prior.count = 1). To express SLE gene expression relative to the control baseline, gene-wise means and standard deviations were estimated from CTRL samples (N=263) on the logCPM scale. SLE samples were then standardized to **z-scores** for each gene. The resulting scaled expression matrix was restricted to SLE samples (N = 226). Molecular subgroups were harmonised to match the original article: “Inflammatory” (N=48), “Healthy-like” (N=66), “Lymphoid” (N=29) and “Interferon” (N=83).

#### Longitudinal cohort (microarray)

Expression matrices were retrieved from GEO (GSE121239). Patients with ≥3 clinical visits were retained. Probes were collapsed to unique gene symbols using platform annotations (first occurrence used when multiple probes mapped to the same gene). To harmonise expression scales within each dataset, gene-wise means and standard deviations from healthy controls were used to z-score all SLE visits. Genes with missing values were removed. Clinical metadata were curated and augmented with time since previous visit, cumulative follow-up, remission status (SLEDAI ≤ 2), and flare risk (≥4-point SLEDAI increase within three months). Expression and clinical data were aligned by visit/sample identifiers, and only SLE visits were used downstream. The final cohort included 358 visits from 84 patients.

### Processing spatial transcriptomic samples

Kidney biopsies of lupus patients were obtained through the Biopsy Biobank Cohort of Indiana ^64^. Formalin-fixed paraffin-embedded tissue was sectioned with a thickness of 5 µm and placed on a regular slide. After staining with hematoxylin and eosin, the slides were imaged with a Keyence BZ-X810 microscope. The 10X Visium assay was performed with the use of CytAssyst equipment, according to the manufacturer’s protocol. Reads were sequenced on an Illumina instrument, and aligned to GRCh38 2020-A reference genome using spaceranger 2.0.1

### Cell type deconvolution in spatial transcriptomic data and glomerular annotation

Seurat anchor method ^65^ was used to deconvolute the transcriptomic signature of each spot on its constituent cell types. Labels were transferred from the single-nucleus from the Human Biomolecular Atlas Program / Kidney Precision Medicine Project atlas ^48^. Transfer scores are interpreted as the proportion of each spot signature associated with the reference cell type. Loupe browser was used to annotate spots as glomerular based on underlying histology, and the expression of NPHS2.

### Comparing cell type annotations in single cell cohorts

Marker genes were defined per cell type using the *sc.tl.rank_genes_groups()* function in Scanpy ^62^ (v1.11.5) in both scRNASeq cohorts, with significance thresholds set to an adjusted p-value<0.001 and log2FC>3. To quantify the consistency of cell-type markers between cohorts, Jaccard indices were computed for all pairwise combinations of cell types based on their respective marker gene sets.

### Compositional analysis of immune cell populations

Compositional profiling of immune cell types was performed for both preprocessed single cell cohorts. For each sample, cell type proportions were computed by counting the number of cells per annotated cell group and normalizing by the total number of cells in the sample.

To account for the compositional nature of proportion data and avoid biases introduced by the unit-sum constraint, centered log-ratio (CLR) transformation was applied. Prior to CLR transformation, zeros in the proportion matrix were addressed using multiplicative replacement, which substitutes zeros with small positive values while preserving the compositional structure^66^.

### Pseudobulk profiles

We generated *pseudobulk* expression profiles by aggregating raw counts at the cell type level within each sample for each cohort from preprocessed scRNASeq cohorts. In the Nehar-Belaid et al. cohort, pediatric (n=44) and adult (n=12) samples were processed separately to account for potential age-related differences in immune cell composition, transcriptional profiles, and disease biology. In Perez et al. cohort all samples (n=261) were used to calculate pseudobulk profiles.

### Differential Gene Expression Analysis in pseudobulk data

Differential gene expression (DGE) analysis was conducted using all *pseudobulk* across both cohorts. As we did in previous analysis, in the Nehar-Belaid et al. cohort, pediatric and adult samples were processed separately to account for potential age-related differences in cell-type specific transcriptional profiles.

Within each cell type, genes were pre-filtered, retaining only those with a minimum of 5 counts in at least one sample and a total count of at least 10 across all samples. DGE was performed using pyDESeq2^67^ (version 0.5.3). Design matrices were constructed using conditions (SLE vs. Healthy), with "Healthy" set as the reference group. Log2 Fold changes were estimated via Wald tests, and shrinkage was applied to improve stability for genes with low dispersion estimates.

Significance was defined as an adjusted *p*-value < 0.05. Genes meeting this threshold were considered differentially expressed. In addition to *p*-values and fold changes, t-statistics derived from the Wald test were extracted for each gene and used in downstream analyses, such as pathway scoring and gene correlation.

### Cross-Cohort Concordance of Differential Expression Signatures

To evaluate the consistency of differential gene expression patterns across cohorts, we computed the Pearson correlation of t-statistics obtained from DESeq2’s Wald test for each pairwise cohort comparison (Perez et al., Nehar-Belaid et al.—adult, and Nehar-Belaid et al.—pediatric). For each annotated cell type, we merged differential expression results across the three cohorts by gene ID and retained only genes common to all datasets. Pairwise correlations of t-statistics were computed separately for each cell type. Pearson correlation coefficients were used as a measure of concordance.

### Cross condition pathway activity inference

We inferred differential pathway activity using PROGENy^20^. For each cohort, *t*-statistics from the differential expression analyses were assembled into a gene-by–cell type matrix (13579 genes in Nehar adults, 13952 genes in Nehar pediatrics and 11434 genes in Perez); missing entries were imputed as zero. Pathway activity scores were computed with the univariate linear model (*ulm*) method in decoupleR ^68^ (v2.12.0). Statistical significance was assessed using the corresponding *p*-values, and pathways with *p* < 0.05 were annotated as enriched in the heatmaps.

### Transcription factor activity inference

We estimated transcription factor (TF) activities from preprocessed, normalized pseudobulk profiles using decoupler-py (v2.1.1). Activities were inferred with the univariate linear model (*ulm*) method and the CollecTRI ^21^ prior-knowledge network as the TF–target regulon.

### Pseudobulk preprocessing to use as input in multicellular factor analysis

Given the sensitivity of matrix factorization to noisy inputs, we enforced conservative QC on *pseudobulk* data before multicellular factor analysis, retaining only high-quality samples and genes to limit technical variability and enhance the robustness and interpretability of the inferred factors. Thus, samples were retained only if they contained a minimum of 10 cells and a total of at least 1,000 counts per cell type. Gene-level quality control was performed using the *filter_by_expr()* function from the decoupler-py package (version 2.0.2). Genes were retained if they met a minimum expression threshold of 10 counts in at least one sample and a cumulative count of at least 15 across all samples. This filtering was applied separately for each cell type within each cohort. To further refine input features, highly variable genes (HVGs) were identified per cell type using Scanpy’s highly_variable_genes() function, specifying cohort-specific batch keys (matching the batch structure used for MOFA grouping). For each cell type, the intersection of HVGs with the genes that passed quality control constituted the final feature set used as input to the MOFA model.

### Multicellular factor analysis

To uncover coordinated transcriptional programs across immune cell types, we applied multicellular factor analysis ^16^ to the joint set of pseudobulk gene-expression profiles. For each cell type and cohort, the model input genes were defined as the intersection of QC-passed genes and highly variable genes; to prevent dominant cell-identity signals from driving the latent structure, we removed the top 25 marker genes per cell type based on differential-expression rankings. The input to the model comprised normalized expression matrices from nine immune cell types. To account for technical batch effects, we used multi-group MOFA framework ^69^, which learns a shared set of factors (i.e., common gene loadings across cell types) while allowing sample-level factor scores to vary across groups. The Nehar-Belaid *et al.* cohort was stratified into pediatric and adult groups, and the Pérez *et al.* cohort was grouped according to the four subcohorts described in the original publication. Model fitting was performed with MOFA2 (mofapy2 v0.7.2) using scale_views=True, likelihoods=’gaussian’, and n_factors=20. Training used fast convergence with dropR²=0.001 and a fixed random seed to ensure reproducibility.

To characterize the biological and clinical relevance of the identified multicellular programs, we associated factor scores with clinical and demographic variables.

First, to globally rank covariates by the proportion of factor variance they explain, we fit a multivariate linear model for each covariate (sex, disease status, age, and ancestry) using all factor scores as predictors, and report the corresponding R².

Subsequently, to map fine-grained clinical associations within SLE patients, we applied a univariate linear model for each (factor, covariate) pair across a panel of 24 variables, including demographics (age, sex, ancestry), disease activity (SLEDAI score), medications (oral corticosteroids, mycophenolate mofetil, methotrexate, hydroxychloroquine), autoantibodies (ANA, anti-dsDNA, anti-Smith, antiphospholipid), and clinical manifestations (lymphopenia, leukopenia, low complement, malar rash, renal involvement, neurologic involvement, arthritis, oral ulcers, serositis, thrombocytopenia, discoid rash, photosensitivity). For each model, a Type II ANOVA F-test was used to derive p-values, eta-squared (η²) to quantify effect size, and the regression t-statistic to capture directionality. P-values were corrected for multiple comparisons across factors per covariate using the Bonferroni method.

### Multicellular networks

For each multicellular program, we quantified per–cell-type activity at the pseudobulk level using the univariate linear model (*ulm*) method implemented in decoupler-py (v2.1.1). Specifically, gene loadings for a given factor and cell type (e.g., B-cell loadings from Factor 1) were used as signed weights to score the corresponding pseudobulk profile for that cell type in each sample. This procedure was repeated for all cell types, yielding, for each factor, an activity matrix with *m* samples (rows) and *n* cell types (columns); in our data *n=9*. Activity scores were then z-score standardized across samples before downstream modelling.

To assess cross–cell-type predictability, we trained ridge regression models (RidgeCV, scikit-learn) to predict the activity of one cell type from the activities of all other cell types. One model was fit per target cell type (total of *n* models). Model performance was internally evaluated using 5-fold cross-validation, reporting the coefficient of determination (R2) on held-out samples for each target. From each fitted model, we extracted the regression coefficients (betas) as a directed measure of the contribution of each predictor cell type to the target. The full procedure was run separately for all samples combined, SLE patients only, and healthy controls (HC) only.

These relationships were summarized as a cell-type–by–cell-type matrix in which entry (X, Y) contains the β coefficient of predictor X in the model for target Y (directed edge X → Y). The model-level R² for each target was retained as a measure of multicellular predictability. To identify condition-specific interactions, we computed the difference in β coefficients between HC and SLE networks. Pairs where the coefficients had opposite signs across conditions, or where the absolute difference was below a magnitude threshold of 0.1, were set to zero (shared).

Remaining pairs were classified as HC-enriched (β_HC > β_SLE) or SLE-enriched (β_SLE > β_HC) and visualized as a cell-type–by–cell-type heatmap. SLE-specific interactions were additionally rendered as a directed graph using, with edge width proportional to the magnitude of the condition difference.

### Interferon-specific cell-cell communication interactions

Cell–cell communication (CCC) was inferred per sample in the preprocessed single-cell cohorts using the LIANA-py package (v1.5.1) ^70,40^, which integrates multiple ligand–receptor inference methods into a consensus interactions score. We restricted the analysis to patients retained for multicellular factor analysis. For each cohort, normalized, log1p-transformed counts were supplied to li.mt.rank_aggregate.by_sample(), grouping cells by annotated cell type and iterating over samples. Using LIANA’s consensus ligand–receptor (LR) resource, we required ligand and receptor expression in ≥10% of cells within the sender and receiver groups (expr_prop = 0.1) and used 100 permutations (n_perms = 100).

Method-specific scores were aggregated with liana’s default magnitude_rank, a rank-based score in which higher values indicate stronger interactions. This yielded, for each sample, LR quadruplets (source cell type – ligand – receptor – target cell type) with an associated score. The resulting per-sample interaction scores were then structured into a multi-view object using li.multi.lrs_to_views(), retaining only LR pairs present in ≥30% of samples (lr_prop = 0.3), views with ≥20 interactions per sample and ≥20 LR pairs, and views covered by ≥100 samples. Missing LR values across samples were filled with zero.

To constrain the CCC landscape to biologically relevant sender–receiver pairs, we leveraged the multicellular networks inferred from a specific factor (Factor 1). Only CCC views whose sender–receiver cell-type pair corresponded to an edge in the respective combined (all-sample) multicellular network were retained.

Finally, for each retained LR quadruplet, we computed the Pearson correlation between its per-sample interaction score and the corresponding MOFA+ factor score across all samples. LR interactions with r > 0.3 were considered factor-associated and carried forward for interpretation.

### Coordinated gene signatures

To do an overrepresentation analysis and for spatial mapping onto tissue, we summarized a program in a single gene signature which aimed to capture genes shared across the most relevant cell types within a program. Guided by the inferred multicellular network, target cell types with cross-validated performance *R*^2^*≥0.8* were deemed coordinated and retained. The final cell-type set per factor was the union of (i) coordinated cell types and (ii) cell types for which the MOFA variance explained exceeded 10%. For factors where the explained variance was restricted to only one cell type (i.e. Factor4 and Factor10) we kept only that cell type.

For each factor, genes were restricted to those with absolute MOFA loading ∣w∣≥0.5 in the selected cell types. A single weight per gene was then obtained by averaging the retained loadings across those cell types. The resulting per-factor weight vectors defined the coordinated gene signatures

### Archetypal definition within each cell type

Archetypal analysis identifies the extreme points of the transcriptional state space within a given cell type. Under the Pareto optimality framework, these extreme points, or archetypes, represent specialist cell states associated with distinct transcriptional tasks. Individual cells are positioned along a continuum between archetypes, reflecting the extent to which they are biased toward one or a combination of these specialized programs^41^.

For each cell type, we subsampled cells belonging to the specific cell type in each single-cell resolution cohort and applied QC filtering (retaining cells with ≥100 detected genes and genes expressed in ≥3 cells). Counts were normalized and log1p-transformed, then scaled to unit variance and zero mean prior to dimensionality reduction. We computed PCA and integrated cohorts using Harmony (Scanpy v1.11.5). The number of retained principal components was selected per cell type based on explained variance: 16 for cDC and cM, 13 for pDC, 12 for T4, 10 for NK and B cells, 9 for proliferating cells, and 7 for ncM and T8.

Archetypes were inferred with ParTIpy (v0.0.1) ^53^ using the harmony embedding. For each cell type, we scanned a specific range of k values and selected the optimal number of archetypes primarily by information criteria, supported by variance explained. The search ranges and resulting optimal k were: pDC (k ∈ {4,…,6}, k=5), cDC (k ∈ {3,…,6}, k=3), cM (k ∈ {4,…,7}, k=6), and k=4 for Prolif (k ∈ {3,…,5}), ncM (k ∈ {3,…,4}), NK (k ∈ {3,…,5}), B (k ∈ {3,…,4}), T4 (k ∈ {4,…,7}), and T8 (k ∈ {3,…,5}).. To assess robustness, we performed a bootstrap stability analysis with 50 resamplings and evaluated the reproducibility of archetype assignments across runs.

To characterize and functionally interpret archetypes, we first computed cell weights for each archetype by using a radial basis function (RBF) kernel on the distances from cells to archetype vertices; the kernel length scale was set to one-half of the median distance from the data centroid to the archetypes. These weights were used for metadata enrichment. Cells were annotated as Healthy or SLE according to patient-level metadata and used to compare archetype enrichments across clinical groups. Second, to associate molecular functions with each archetype, we constructed archetype-specific gene expression profiles by weighting the z-scored expression matrix with the corresponding cell weights, enabling comparison of gene expression patterns across archetypes.

### Association of archetypes to multicellular program

We investigated whether patients with higher or lower program scores preferentially occupy cell states closer to specific archetypes, enabling cross-condition comparison in a shared latent space. For each patient, single-cell gene expression profiles were reduced to a single patient-level summary by averaging across all cells from that patient. This mean vector provides a first-order description of the patient’s overall cell-state distribution and ensures that downstream analyses use one independent observation per patient. Prior to aggregation, we verified that factor loadings and patient-level covariates (batch and demographic variables) were constant within each patient.

Because patient mean profiles can lie slightly outside the convex region defined by the archetypes, we projected each patient mean onto the convex hull spanned by the archetypes. This step ensures that subsequent proximity calculations are interpreted relative to the archetypal task space itself and avoids inflating distances due to out-of-hull points. The deviation between the original patient mean and its projection was retained as a diagnostic measure of how far a patient lies outside the archetype simplex and could be used as an additional nuisance covariate when needed.

We then quantified patient proximity to each archetype by computing Euclidean distances between the projected patient mean and each archetype, yielding one distance per patient–archetype pair. Distances were analyzed on their original scale by default; in sensitivity analyses, we also rank-transformed distances to reduce sensitivity to differences in scale across conditions.

To test for association between factor loadings and archetype proximity, we fit separate patient-level linear regression models for each archetype, with the patient’s program loading as the outcome and the distance to that archetype as the main predictor. Models were estimated using ordinary least squares, and statistical inference used heteroskedasticity-consistent (HC3) standard errors to account for potential non-constant variance. Associations were interpreted such that a negative distance coefficient indicates that patients closer to a given archetype tend to have higher factor loadings, whereas a positive coefficient indicates that patients closer to that archetype tend to have lower factor loadings. Multiple testing across archetypes was controlled using either Bonferroni or Benjamini–Hochberg false discovery rate correction, depending on the analysis scope. As a robustness check, we additionally assessed significance using patient-level permutation tests: factor loadings were permuted across patients when no covariates were included, and Freedman–Lane residual permutation was used when covariates were included to preserve the covariate structure.

### Cytokine enrichment of archetypes

To characterize the cytokine signaling landscape of each archetype, we estimated cytokine activity scores using the Human Cytokine Response Atlas (huCIRA)^44^, a prior knowledge network that provides transcriptional response signatures for 90 cytokines across 24 immune cell types, defined by log fold changes of target genes upon cytokine stimulation. The network was formatted as a weighted bipartite graph where edges connect cytokine–cell type source nodes to target genes, weighted by log fold change.

Archetype gene expression profiles were represented as z-score-scaled mean expression vectors. Profiles from all 9 cell types (38 archetypes total) were concatenated into a unified matrix and cytokine activity scores were estimated using the univariate linear model method implemented in decoupler-py (v2.1.1). To ensure specificity, each archetype’s scores were restricted to cytokine signatures pertaining to its own cell type (e.g., classical monocyte archetypes matched to CD14+ monocyte perturbation signatures).

To assess the similarity of SLE-associated archetypes in cytokine activity space, we constructed a cytokine activity matrix restricted to SLE archetypes and to cytokines significant in at least one SLE archetype (adjusted *P* < 0.05). Pairwise cosine similarity was computed between archetype vectors and visualised as a clustered heatmap using Ward’s linkage.

Then, for cytokines present in both SLE- and HC-associated archetype groups, the mean ULM score was computed across each group and the difference (Δ = mean_SLE − mean_HC) was used as an effect size. Statistical significance was assessed per cytokine using a two-sided Wilcoxon rank-sum test, with BH correction applied. Cytokines with FDR < 0.05 were retained.

### Linking archetype transcriptional signatures to circulating protein biomarkers

To assess whether genes upregulated in SLE-associated archetypes encode proteins detectable in peripheral blood, we integrated archetype expression profiles with plasma proteomics data from the Disease Blood Atlas^45^. For each archetype, gene expression was represented as a z-score computed per gene across all archetypes of the same cell type, such that positive values indicate relative upregulation compared to the cross-archetype mean. A gene was considered upregulated in an archetype if its z-score exceeded 0.2, a threshold chosen to retain genes expressed above the cross-archetype mean. In addition, proteins significantly elevated in SLE patient plasma relative to healthy donors (adjusted P < 0.05) were retrieved from the Disease Blood Atlas (n = 432), and their assay identifiers were matched to gene names in the archetype expression matrix. To identify proteins with selective transcriptional support in disease states, we retained only those upregulated in at least one SLE-associated archetype and in no HC-associated archetype. The resulting 47 proteins were visualized as a heatmap of z-scored expression across SLE archetypes, with hierarchical clustering applied to both rows and columns.

### Mapping multicellular programs onto bulk-transcriptomic cohorts

To assess the activity of multicellular programs in external bulk transcriptomic data, we mapped previously inferred latent factors onto an independent cohort using a weighted scoring approach.

First, factor loadings (gene weights per cell type and factor) were extracted from the model and reshaped into a regulatory network where each edge links a gene (target) to a factor-cell type pair (source) with an associated weight. This network was used as input to the Univariate Linear Model (ulm) implemented in decoupler-py (v2.1.1) ^68^, which estimates enrichment scores (activity scores) for each factor-celltype combination in each sample. The result was a mapping matrix of dimension samples × factor-celltype combinations, with corresponding significance values.

To compute a factor activity score per sample and account for cell type-specific contributions to each multicellular program, we first extracted the proportion of variance explained (R²) for each factor across all groups. These R² values were used as weights to compute a weighted average of factor contributions across cell types for each sample in the mapping matrix.

### Cox proportional hazards analysis of immune programs and flare risk

To assess whether multicellular programs scores at the time of visit were associated with the risk of subsequent disease flare, we applied Cox proportional hazards models in a longitudinal cohort of adult SLE patients. For each patient visit, we mapped cell-type program activities (see bulk mapping). To obtain a single factor score per visit, activities were aggregated as a weighted mean across cell types, where weights were defined as the sum of variance explained (R²) by each factor in each cell type. Analyses were restricted to visits during clinical remission. Visits were labelled as Short-Term if the patient experienced a flare within the following 3 or 6 months, and as Long-Term if no flare occurred within that window, the last recorded SLEDAI score was below 2, and cumulative follow-up exceeded 90 days. Only visits falling into one of these two categories were retained, yielding a paired comparison between imminent flare and sustained remission. Survival was defined as time (in days) from the index visit to the next scheduled visit, with flare status as the event indicator. We fitted univariate Cox models for four biologically annotated factors: IFN, NF-κB, T8 cytotoxic, and T4 naive/effector program. To evaluate the independent contribution of each program beyond the dominant IFN and NF-κB signals, pairwise multivariate Cox models were fitted (e.g., NFkB + IFN, T8 cytotoxic + IFN). Model performance was assessed with the concordance index (C-index). Likelihood ratio tests (LRT) were used to compare nested models and determine whether adding a factor significantly improved predictive fit beyond the base model. All analyses were performed independently for the 3-month and 6-month prediction windows using the survival R package ^71^(v3.5.8).

### Mapping multicellular programs onto spatial transcriptomic cohort

We projected the previously defined coordinated gene signatures (one per factor) onto 10x Visium slides to obtain per-spot enrichment scores. Visium count matrices were variance-stabilised and scaled prior to scoring. Enrichment was computed with decoupleR (v2.12.0) using the univariate linear model (ULM) method, applied separately to each factor–signature.

For visualisation, we focused on immune-activation programs (Factors 1, 2, and 4). For each slide, negative enrichment values were set to zero to obtain non-negative, relative abundances across the three factors; the mean of these three non-negative scores was used to define the pie-chart intensity.

To quantify factor enrichment in glomeruli, we first selected glomerulus-annotated spots across slides and removed spots with predicted immune (IMM) cell-type composition equal to 0 (level-1 deconvolution). Remaining spots were labelled SLE or Healthy from patient-level metadata. For each factor, we fit a linear model of spot-level enrichment versus condition (SLE vs Healthy) and extracted the *t*-statistic and *P*-value to assess whether enrichment was higher in SLE glomeruli.

### Spatial co-localization between immune programs and kidney cell types abundances

To examine whether MOFA+ immune programs are spatially organised within the renal tissue, we quantified the co-localisation between factor activity scores and cell-type deconvolution estimates across spots in visium spatial transcriptomics slides.

Spatial weight matrix. For each tissue section, a k-nearest neighbour (k = 6) spatial weight matrix W was constructed from spot coordinates, where W_{ij} = 1 if spot j is among the 6 nearest neighbours of spot i, and 0 otherwise.

Co-localisation between a factor score vector x and a cell-type abundance vector y was quantified using the bivariate global Moran’s I. Statistical significance was assessed by a permutation test (n = 100 permutations) in which the y vector was randomly shuffled to build the null distribution. Two-tailed p-values were computed using the Davison & Hinkley correction, setting a minimum achievable p-value of 1/101. This was computed for all combinations of 4 immune factors (IFN, NF-κB, T8 cytotoxic, T4 naive/effector) and cell-type deconvolution scores in each slide, separately for SLE and healthy samples.

Per-slide Moran’s I values were averaged across all slides within each condition (SLE or healthy). To combine evidence across independent samples, p-values were aggregated using Fisher’s combined probability test, distributed as χ² with 2k degrees of freedom (k = number of samples). The resulting combined p-values were corrected for multiple testing across all (cell type × factor × condition) combinations using the BH method. Associations with FDR < 0.05 and |mean Moran’s I| ≥ 0.10 were considered significant.

### Glomerular cell-type composition and neighbourhood context

To quantify the spatial distribution of cell-type compositions across slides, we applied MISTy (1.16.0) ^51^ to 10x Visium data with manually annotated glomerular spots. For each slide, per-spot cell-type compositions were inferred with Seurat using first-level annotations. Compositions <0.05 were set to 0, and cell types with zero variance across spots were removed. Remaining compositions were centred log-ratio (CLR) transformed after adding a pseudocount of 1×10^−3^^1^ to avoid infinities.

Two MISTy views were defined. The *intraview* comprised the per-spot CLR-transformed compositions. The *paraview* captured local neighbourhood signal: spot coordinates were retrieved from the Seurat object, pairwise Euclidean distances between spots were computed, and a data-driven nearest-neighbour spacing was estimated from the distance spectrum. The paraview interaction radius *r* was set to twice this spacing (*l=2*) to include second-neighbour interactions. Paraview features were computed as distance-weighted means of neighbouring spots within *r* (weights decreasing with distance, following the original MISTy formulation). After constructing the paraview, slides were masked to retain glomerular spots and neighbours within two-spot distance.

Multi-view models were fitted with MISTy using a Random Forest base learner, combining the intrinsic and paraviews to estimate view contributions and feature importances. Results were then aggregated by condition (SLE vs Healthy) across slides to enable condition-level comparisons.

## Data availability

The single-cell RNASeq datasets used in this study are available at GSE174188 and GSE135779. The bulk transcriptomic longitudinal cohort is available under GSE121239. Bulk transcriptomic data from the PRECISESADS cohort is available under request, as authors mentioned in the original manuscript^3^. Spatial-resolved datasets from kidney biopsies will be provided in public repositories upon publication. The processed transcriptomic data are available on Zenodo (DOI: 10.5281/zenodo.19811831)

## Code availability

All code associated with this publication is available at https://github.com/saezlab/metalupus_pub

## 5. Acknowledgements

J.L.B. acknowledges the support of the Galician Government through the fellowship ED481B_072. P.S.L.S. has received funding from the Deutsche Forschungsgemeinschaft under grant agreement SPP 2395. L.Z. is supported by the Multi-dimensionAI project (CZS-Project number P2022-08-010) funded by the Carl-Zeiss-Stiftung. This work was supported by the Innovative Medicines Initiative joint undertaking Grant # GA115565 (M.E.A.R), by Swedish Research Council grant: 01000-2022 (M.E.A.R) and by grant PID2024-156297OB-I00 funded by MICIU /AEI /10.13039/501100011033/ (ERDF, EU) (P.C.S). The results presented here are in part based upon data generated by the Kidney Precision Medicine Project (KPMP). The KPMP is supported by the National Institute of Diabetes and Digestive and Kidney Diseases (NIDDK) through the following grants: U01DK133081, U01DK133091, U01DK133092, U01DK133093, U01DK133095, U01DK133097, U01DK114866, U01DK114908, U01DK133090, U01DK133113, U01DK133766, U01DK133768, U01DK114907, U01DK114920, U01DK114923, U01DK114933, U24DK114886, UH3DK114926, UH3DK114861, UH3DK114915, and UH3DK114937. The authors gratefully acknowledge the data storage service SDS@hd supported by the Ministry of Science, Research and the Arts Baden-Württemberg (MWK) and the German Research Foundation (DFG) through grant INST 35/1803-1 FUGG and INST 35/1804-1 LAGG. We gratefully acknowledge support by the state of Baden-Württemberg through bwHPC and the German Research Foundation (DFG) through grant INST 35/1597-1 FUGG. We thank Leonie Küchenhoff, Daniel Guerrero Romero and Ines Rivero for critical feedback on the manuscript. Importantly, we acknowledge all data authors that are cited in this study.

## 7. Authors contributions

J.L.B.: data curation, formal analysis, investigation, methodology, software, writing–original draft preparation. P.S.L.S, L.Z. and R.M.F: formal analysis, validation, writing–revision. D.T.D., P.C.S. and J.T.: investigation. M.E.A.R and M.T.E.: supervision, resources. R.O.R.F: supervision, project administration, writing–revision. J.S.R.: supervision, project administration, funding acquisition, writing–revision.

## Supplementary Materials

### Supplementary Files

#### Supplementary File 1

**Sup. Table 1:** Explained variance of multicellular programs

**Sup. Table 2:** Cell-cell communication interactions associated with IFN program. Ligand-receptor (LR) pairs with a Pearson correlation coefficient > 0.3 between their communication activity score (estimated by LIANA+) and IFN program of the multicellular factor model.

**Sup. Table 3:** Statistics for the archetype association to the IFN program

**Sup. Table 4:** Cytokine footprint activity differences between SLE and healthy archetypes

**Sup. Table 5:** Cox proportional hazards models using longitudinal cohort.

**Sup. Table 6:** Performance metrics from MISTy models to study spatial co-localization of cell-types between healthy and SLE slides

### Supplementary Figures

**Supplementary Figure S1.**
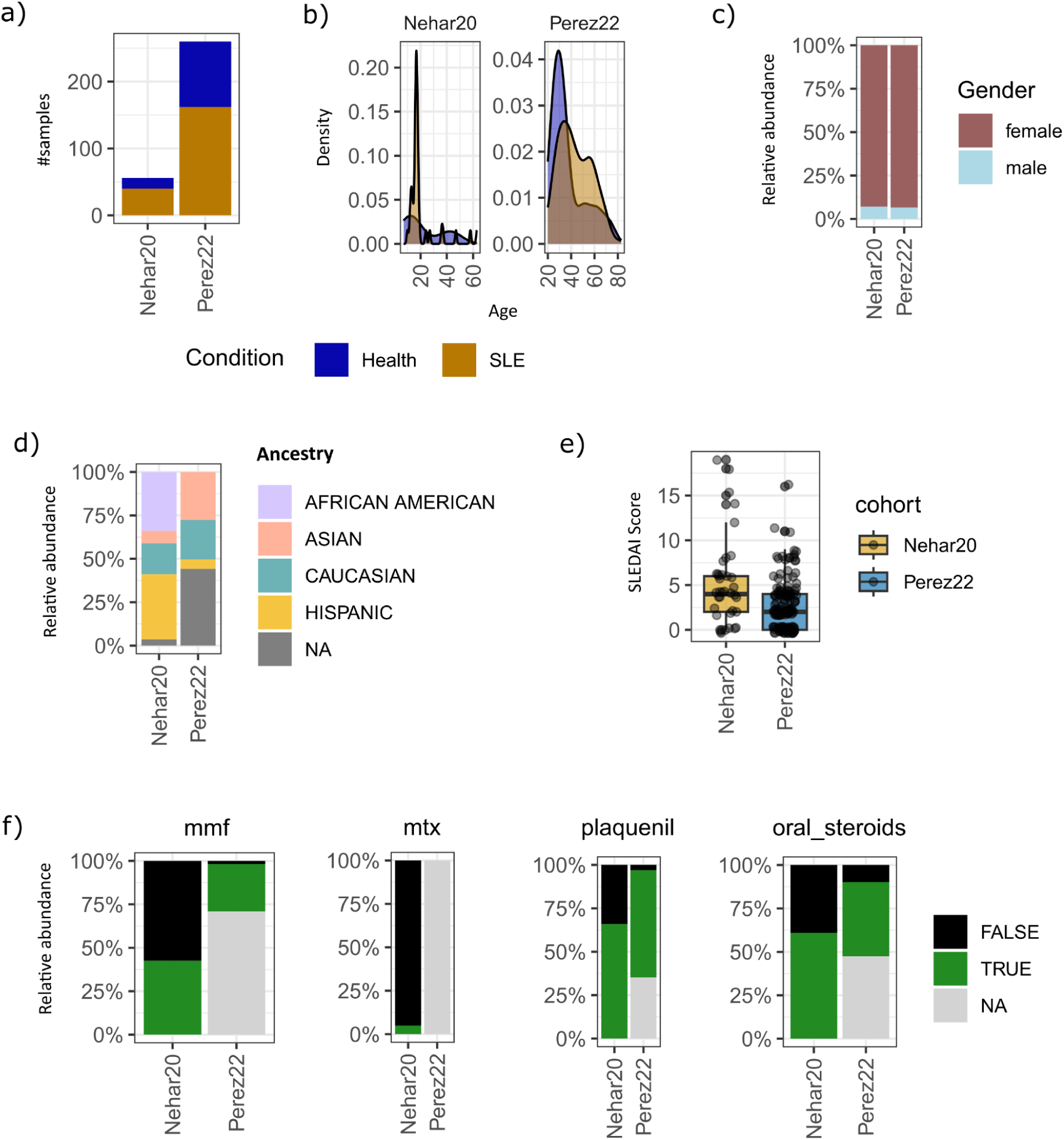
Patient-level clinical and demographic characteristics. (A) Number of samples per cohort (Nehar-Belaid 2020, Pérez 2022) stratified by condition (Healthy, SLE). (B) Age distribution of participants by cohort and condition. (C) Sex distribution across cohorts. (D) Self-reported ancestry composition for individuals with available data. (E) Distribution of disease activity scores (SLEDAI) in SLE patients by cohort. (F) Proportion of SLE patients receiving immunomodulatory treatments, including mycophenolate mofetil (MMF), methotrexate (MTX), hydroxychloroquine (Plaquenil), and oral corticosteroids. Grey indicates missing data.

**Supplementary Figure S2.**
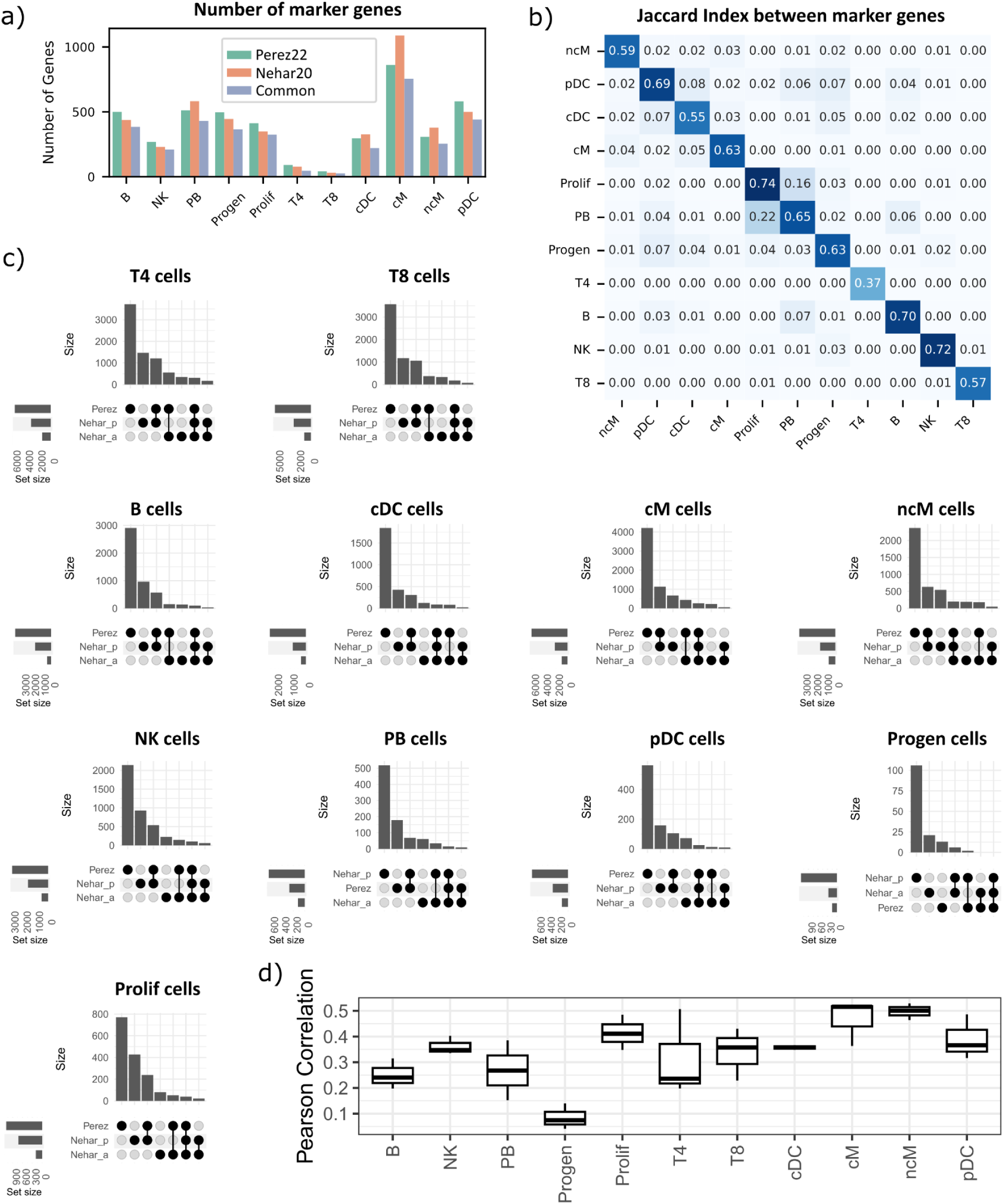
Cross-cohort consistency of cell-type marker and differential expression signatures. (A) Number of marker genes identified per cell type (logFC > 3; P < 0.001). Bars represent markers specific to each cohort (green and orange) and those shared between cohorts (blue). (B) Pairwise Jaccard index heatmap showing the similarity of marker gene sets across all immune cell types between cohorts. (C) UpSet plots displaying intersections of differentially expressed genes (adjusted P < 0.05) identified in SLE versus Healthy comparisons across cohorts. The Nehar-Belaid dataset was divided into pediatric and adult subgroups for independent analysis. (D) Distribution of Pearson correlation coefficients (r) comparing t-statistics from differential expression analyses across cohorts. Each boxplot summarizes three pairwise comparisons (Pérez vs Nehar-pediatric, Pérez vs Nehar-adult, and Nehar-pediatric vs Nehar-adult).

**Supplementary Figure S3.**
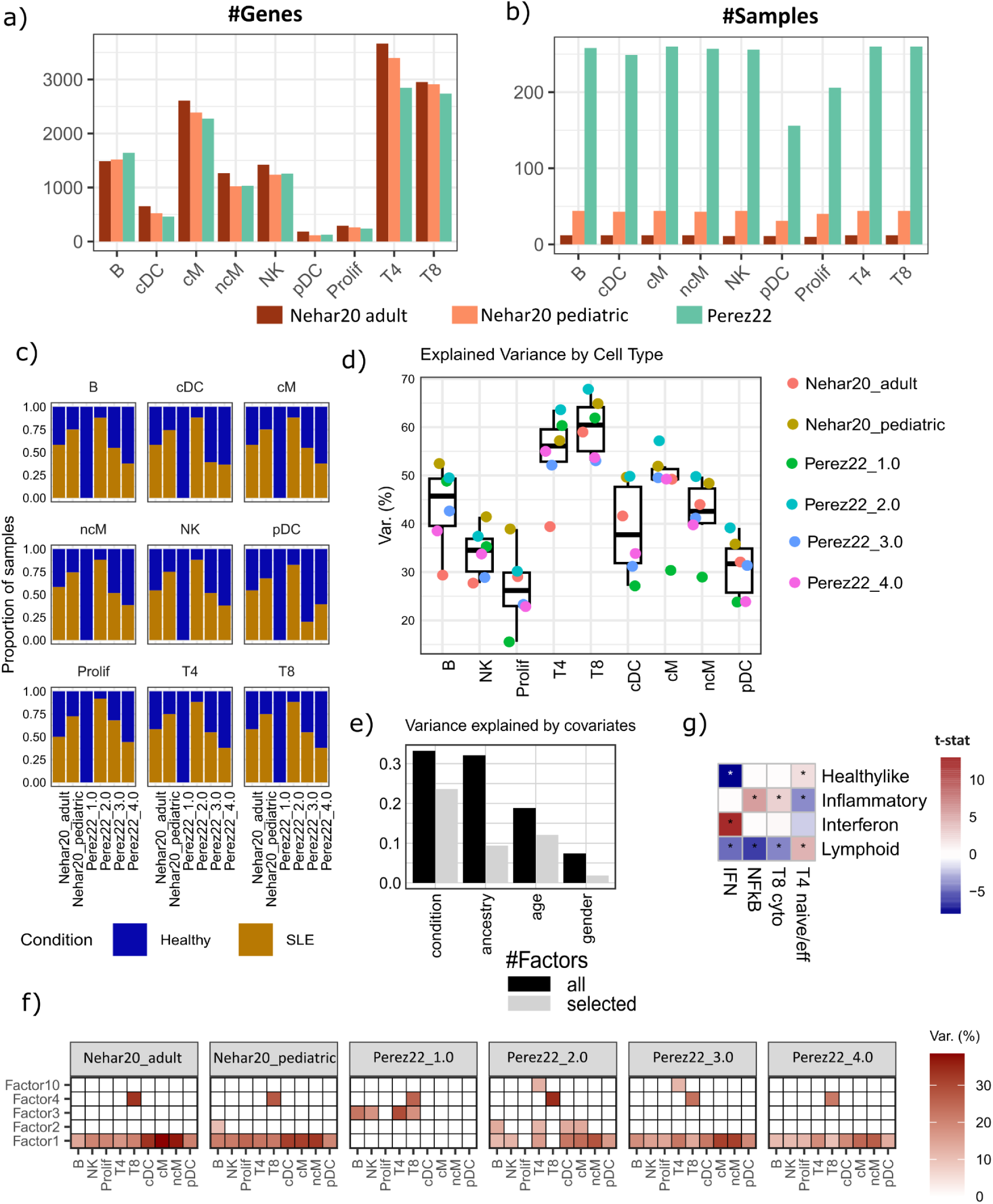
Overview of input data and explained variance of the multicellular factor model. (A) Number of genes per cell type and cohort used as input for the multicellular factor model. (B) Number of samples per cell type and cohort used as input for the multicellular factor model. (C) Proportion of healthy and SLE samples across all cell types and cohorts used as input for the multicellular factor model. (D) Mean explained variance (*R²*) per cell type across cohorts after model training. (E) Variance explained by clinical and demographic covariates (condition, ancestry, age, gender, cohort) when considering all latent factors (*n* = 19) or the selected subset (*n* = 4), estimated via multivariate linear regression. Factor 3 was excluded, as it predominantly captured variation associated with healthy samples. (F) Explained variance (*R²*) across cohorts, cell types, and factors. Only factors explaining ≥10% of the variance in at least one cohort or cell type are shown; others were excluded. For visualization, *R²* values below 10% were set to 0. (G) Mapping multicellular programs onto an external cohort (PRECISESADS, n = 226 SLE patients) and enrichment of these programs in molecular groups defined in the original article (Interferon, Inflammatory, Lymphoid and Healthy-like). Tiles display t-statistics from linear models, with color indicating directionality and asterisks denoting significance (adjusted *P val* < 0.05)

**Supplementary Figure S4.**
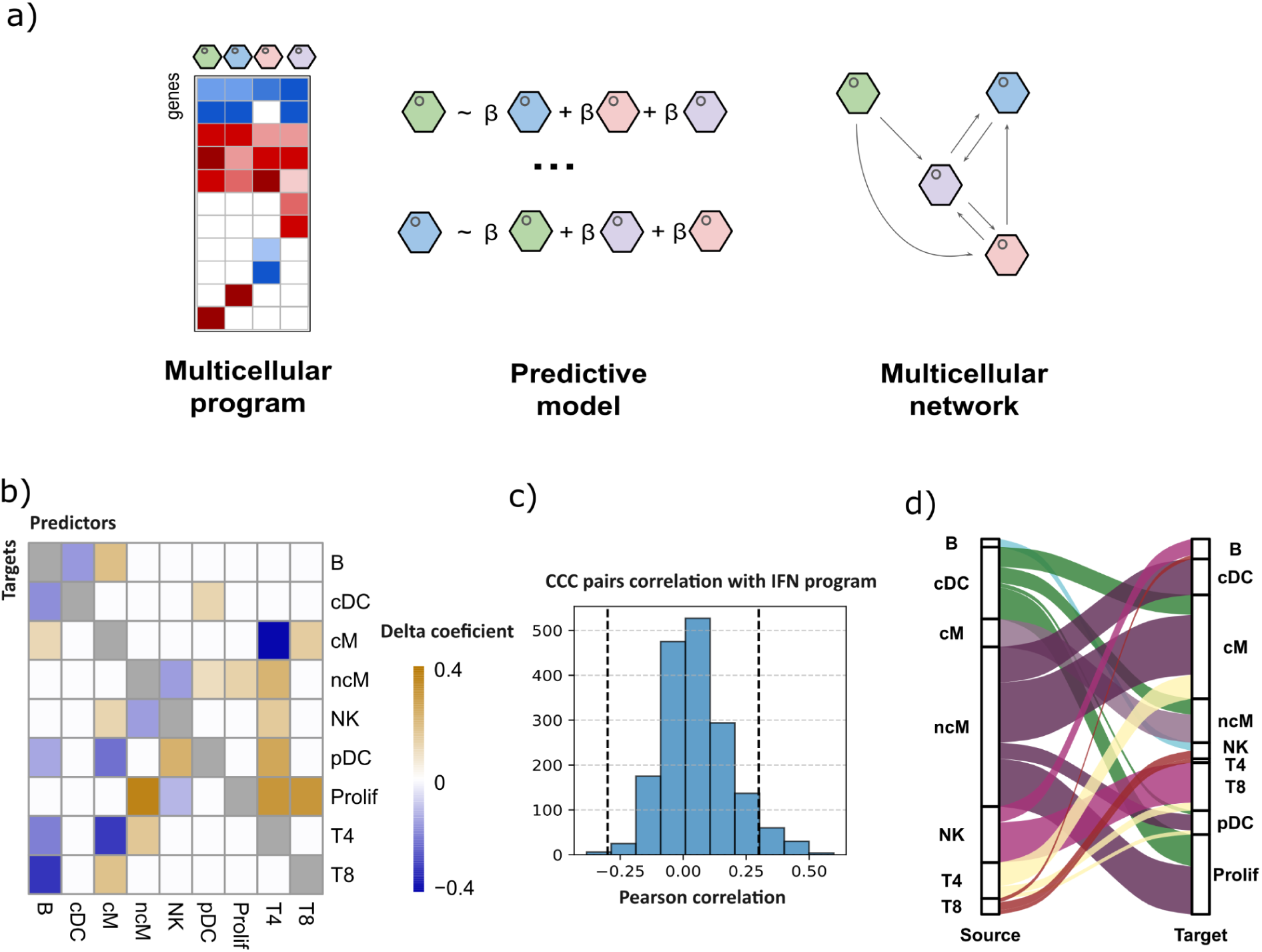
Multicellular coordination and cell-cell communication driven by IFN program. (A) Schematic overview of the framework used to infer multicellular coordination from pseudobulk transcriptomic profiles. Gene-level variation across immune cell types is decomposed into latent factors that capture shared multicellular programs. Intercellular coordination is quantified by predictive modeling of factor loadings across cell types, followed by inference of ligand–receptor (LR) communication links constrained by the resulting co-expression network. (B) Difference in cell-type dependencies within the IFN program between SLE and healthy samples. Each cell shows the difference in ridge regression coefficients (SLE − healthy) for a given predictor (column) - target (row) pair. Coefficients below a magnitude threshold of 0.1 in either condition were set to zero, and pairs where both conditions shared the same sign were excluded. Orange indicates dependencies stronger in SLE; blue indicates dependencies stronger in healthy networks. Diagonal elements (self-predictions) are shown in gray. (C) Distribution of pairwise Pearson correlation coefficients (*r*) between Factor 1 loadings and ligand–receptor (LR) interaction activities inferred using LIANA (see Methods). The dashed vertical line denotes |*r*| = 0.3. CCC interactions were constrained to those identified within the merged multicellular network (Healthy + SLE). (D) Summary of significant ligand–receptor (LR)–mediated cell–cell communication (CCC) events along the *Factor 1* axis. Significant interactions were identified based on Pearson correlation (r > 0.3, *P* < 0.05) between LR interaction scores and *Factor 1* loadings across individuals. Each interaction reflects the co-expression of a ligand in one cell type and its receptor in another, calculated from mean expression values at the pseudobulk level. The chord diagram depicts the number of significant LR interactions between source and target cell types.

**Supplementary Figure S5.**
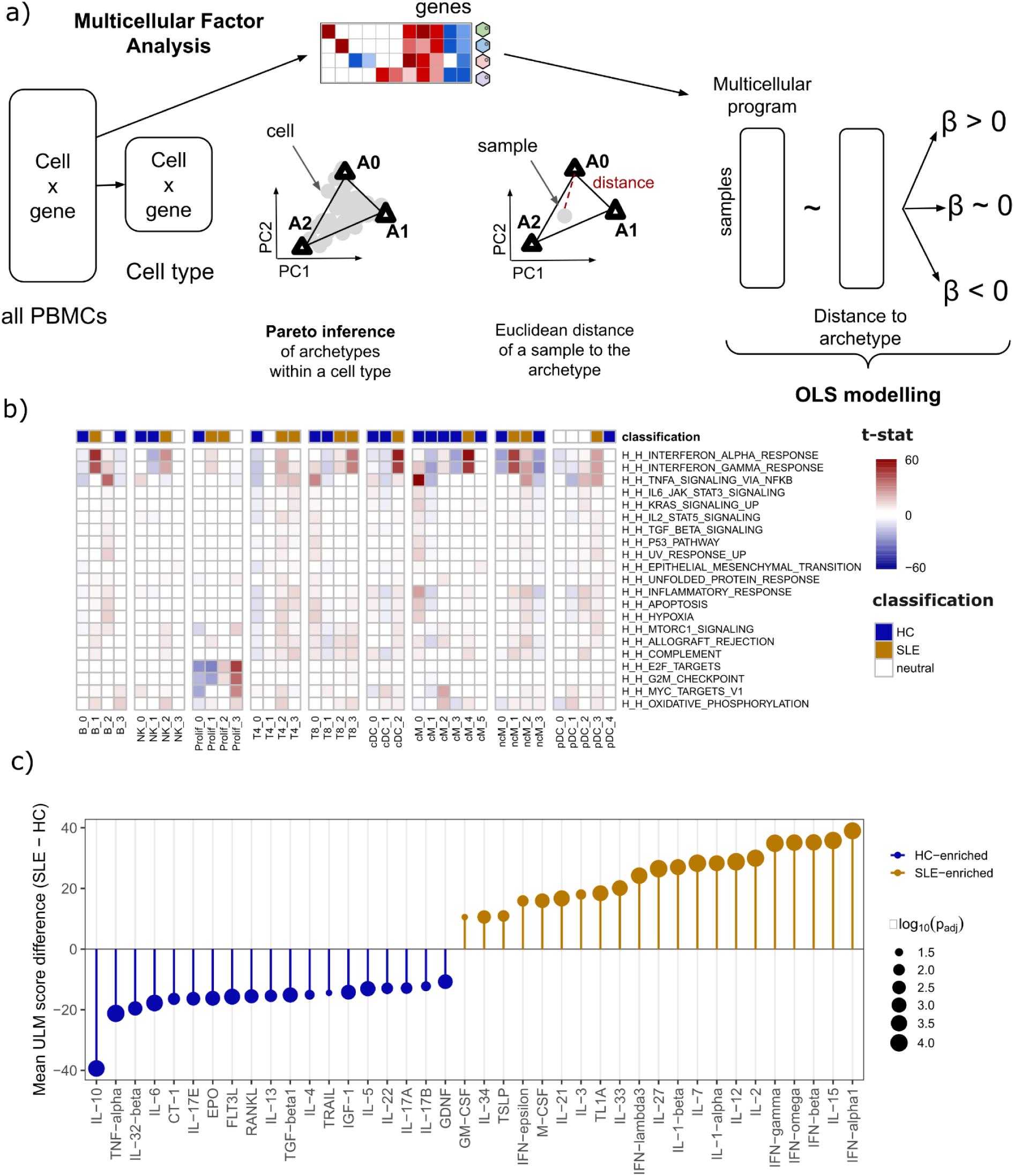
Archetypal analysis. (A) Overview of the methodology to link multicellular programs to archetypal cell states across patients. Within each cell type, single-cell expression profiles are used to identify archetypes using Pareto task inference, and each patient summary is projected onto the archetype simplex. Euclidean distances from each patient to each archetype are then computed. Finally, sample multicellular program scores (from IFN program) are modeled as a function of distance to each archetype using ordinary least squares. Negative regression coefficient indicates that higher program loading is associated with closer proximity to the corresponding archetype, whereas a positive coefficient indicates association with greater distance. (B) Hallmarks enrichment across all archetypes. Heatmap of ULM enrichment scores (t-values) for most enriched Hallmarks across all archetypes (columns), grouped by cell type. Only Hallmarks–archetype pairs with adjusted P < 10⁻⁶ are shown (non-significant entries set to zero). Columns are ordered by cell type; vertical gaps separate cell types. Top annotation indicates archetype classification: SLE-associated (orange), Healthy-associated (blue), or neutral (gray), based on the regression of archetype proximity against IFN program scores (FDR < 0.05). Rows are hierarchically clustered. Color scale represents t-values: red, positive enrichment; blue, negative enrichment; white, non-significant or zero. (C) Differential cytokine enrichment between SLE- and HC-associated archetypes. For each cytokine, the mean ULM enrichment score was computed separately across SLE-associated and HC-associated archetypes, and the difference (SLE minus HC) is shown on the y-axis. Significance was assessed by Wilcoxon rank-sum test with Benjamini–Hochberg correction; top 40 most significant cytokines with adjusted P < 0.05 are displayed. Point size encodes −log₁₀(adjusted P-value). Orange, SLE-enriched cytokines; blue, HC-enriched cytokines.

**Supplementary Figure S6.**
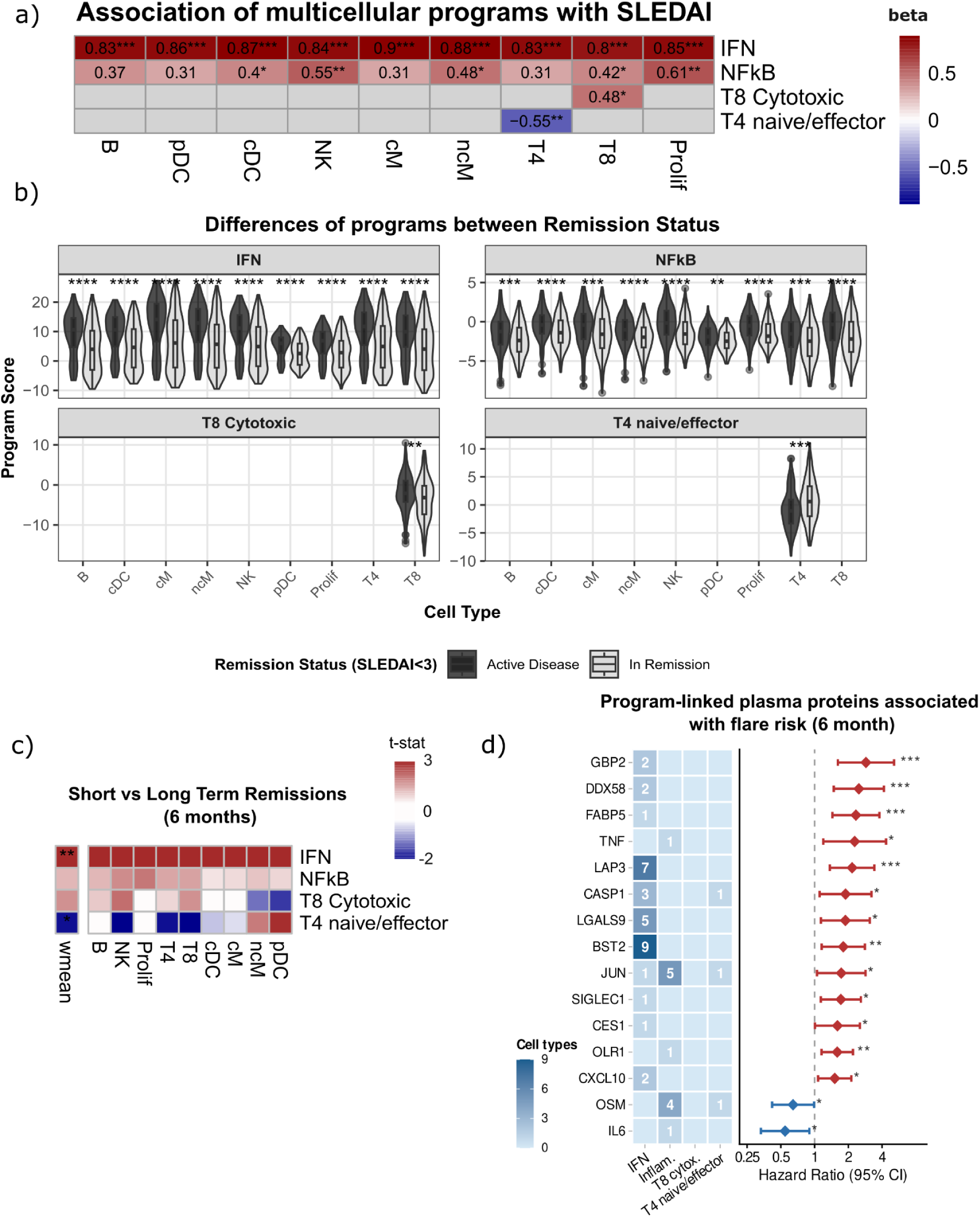
Flare prediction. (A) Heatmap showing the association between multicellular program scores and SLEDAI in the longitudinal adult SLE cohort. Each cell displays the standardized regression coefficient (β) from a linear mixed-effects model. Rows correspond to the four retained multicellular programs and columns to immune cell types. Grey cells indicate program–cell-type combinations excluded from the model due to cell-type specificity of the program. Asterisks denote statistical significance after Benjamini–Hochberg correction: *p < 0.05, **p < 0.01, ***p < 0.001. (B) Violin plots showing the distribution of multicellular program scores across cell types, stratified by active (SLEDAI ≥ 3) vs. remission disease (SLEDAI < 3). Statistical comparisons between groups were performed using the Wilcoxon rank-sum test; significance levels are shown above each comparison (ns: not significant, *p < 0.05, **p < 0.01, ***p < 0.001). (C) Heatmaps showing differential immune program activity between remission visits that preceded a clinical flare (short-term remission) and those with sustained disease control (long-term remission), at a 6-month prediction time frame. For each program–cell type combination, differences were assessed by two-sample t-test; colour represents the t-statistic, with positive values indicating higher program activity in short-term (pre-flare) visits. The top row ("wmean") summarises a weighted mean across cell types, where weights are proportional to the variance explained (R²) by each factor in each cell type as estimated by the multicellular factor model. P-values were corrected for multiple comparisons using the Holm procedure; asterisks denote significance: *p < 0.05, **p < 0.01, ***p < 0.001. (D) Combined plots showing genes significantly associated with flare risk at a 6-month prediction time frame, that overlap with top gene signatures of the multicellular programs and were detected in blood proteomic cohort upregulated in SLE patients. Left panel (heatmap): for each gene, the number of cell types in which it constitutes a top gene of the respective program (IFN, NFkB, T8 Cytotoxic, T4 naive/effector). Right panel (forest plot): hazard ratio (HR, 95% CI) from univariate Cox proportional hazards models, estimating the association between gene expression and time to flare. Proteins are ordered by HR. Red indicates proteins associated with increased flare risk (HR ≥ 1, p < 0.05); blue indicates protective associations (HR < 1, p < 0.05); grey indicates non-significant associations. Asterisks denote significance: *p < 0.05, **p < 0.01, ***p < 0.001.

**Supplementary Figure S7.**
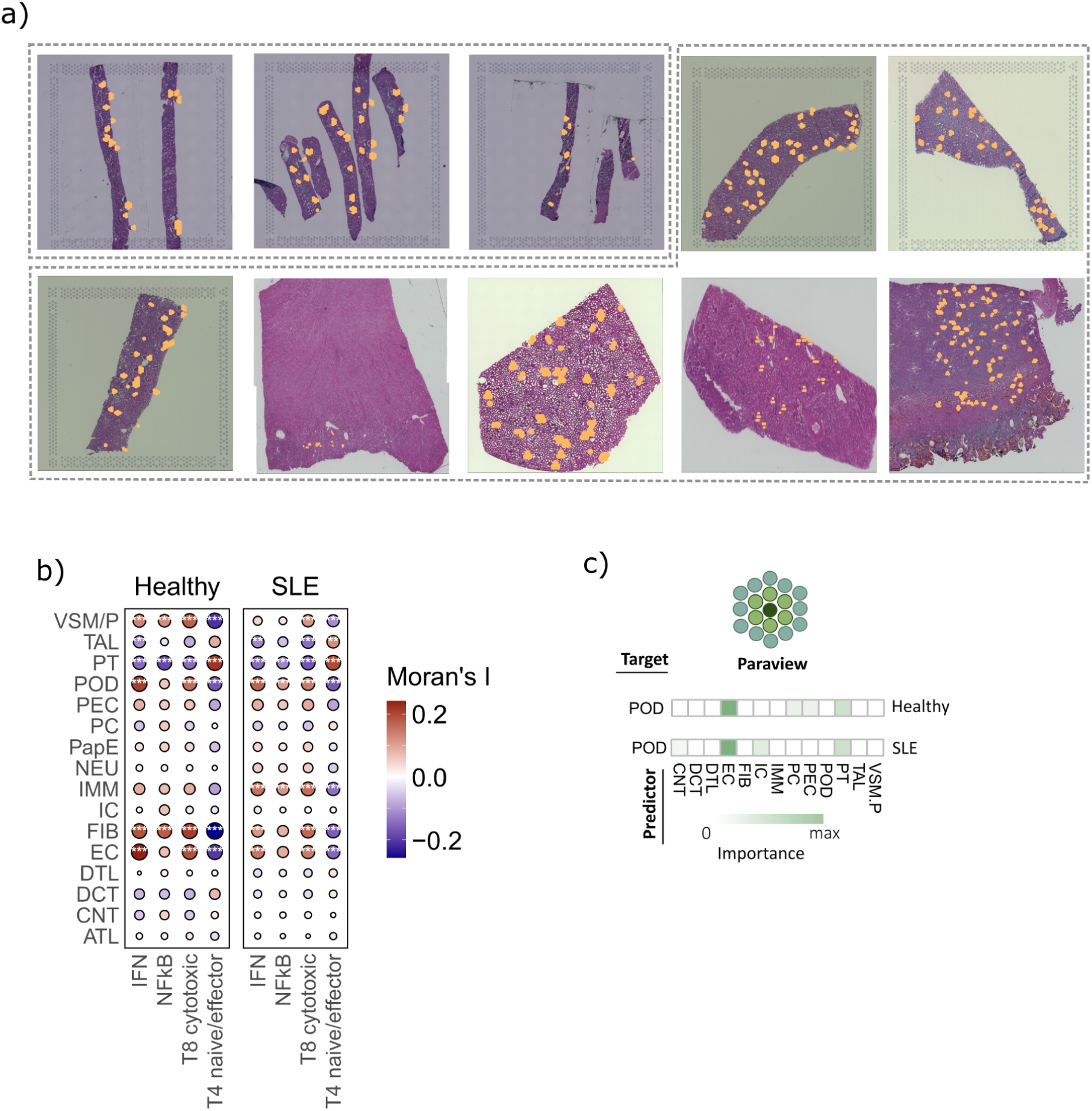
Spatial analysis of cell type composition in kidney samples. (A) Visium 10x spatial transcriptomics slides from SLE (top row, first three) and healthy control kidneys (remaining slides) included in the analysis. Orange dots denote spatial spots annotated as glomeruli. (B) Spatial correlation between multicellular program activities and predicted cell-type abundances between conditions, quantified using bivariate Moran’s I. Moran’s I values were averaged across slides per condition (SLE, Healthy), and statistical significance was assessed by combining per-sample permutation-derived p-values using Fisher’s method, followed by Benjamini-Hochberg FDR correction. Dot size represents the magnitude of spatial association (|Moran’s I|); color indicates directionality (red = positive, blue = negative). Asterisks denote associations that are simultaneously statistically significant (FDR < 0.05) and exceed a minimum effect size (|Moran’s I| ≥ 0.10). (C) Variable importance from the paraview model for predicting key glomerular cell types (endothelial cells, EC; podocytes, POD). Higher importance values indicate stronger spatial influence from neighboring regions.

## References

1. Goldblatt, F. & O’Neill, S. G. Clinical aspects of autoimmune rheumatic diseases. Lancet Lond. Engl. 382, 797–808 (2013).

2. Allen, M. E., Rus, V. & Szeto, G. L. Leveraging Heterogeneity in Systemic Lupus Erythematosus for New Therapies. Trends Mol. Med. 27, 152–171 (2021).

3. Barturen, G. et al. Integrative Analysis Reveals a Molecular Stratification of Systemic Autoimmune Diseases. Arthritis Rheumatol. 73, 1073–1085 (2021).

4. Sandling, J. K. et al. Molecular pathways in patients with systemic lupus erythematosus revealed by gene-centred DNA sequencing. Ann. Rheum. Dis. 80, 109–117 (2021).

5. Kato, H. & Kahlenberg, J. M. Emerging biologic therapies for systemic lupus erythematosus. Curr. Opin. Rheumatol. 36, 169–175 (2024).

6. Toro-Domínguez, D. et al. Stratification of Systemic Lupus Erythematosus Patients Into Three Groups of Disease Activity Progression According to Longitudinal Gene Expression. Arthritis Rheumatol. 70, 2025–2035 (2018).

7. Hubbard, E. L. et al. Analysis of transcriptomic features reveals molecular endotypes of SLE with clinical implications. Genome Med. 15, 84 (2023).

8. Parikh, S. V. et al. Molecular profiling of kidney compartments from serial biopsies differentiate treatment responders from non-responders in lupus nephritis. Kidney Int. 102, 845–865 (2022).

9. Perez, R. K. et al. Single-cell RNA-seq reveals cell type-specific molecular and genetic associations to lupus. Science 376, eabf1970 (2022).

10. Nehar-Belaid, D. et al. Mapping systemic lupus erythematosus heterogeneity at the single-cell level. Nat. Immunol. 21, 1094–1106 (2020).

11. Nakano, M. et al. Distinct transcriptome architectures underlying lupus establishment and exacerbation. Cell 185, 3375–3389.e21 (2022).

12. Gurajala, S. et al. A population-scale atlas of blood and tissue in lupus nephritis. 2025.08.11.669754 Preprint at 10.1101/2025.08.11.669754 (2025).

13. Duan, M. et al. SLECA: a single-cell atlas of systemic lupus erythematosus enabling rare cell discovery using graph transformer. 2026.02.11.705246 Preprint at 10.64898/2026.02.11.705246 (2026).

14. Forlin, R., James, A. & Brodin, P. Making human immune systems more interpretable through systems immunology. Trends Immunol. 44, 577–584 (2023).

15. Jerby-Arnon, L. & Regev, A. DIALOGUE maps multicellular programs in tissue from single-cell or spatial transcriptomics data. Nat. Biotechnol. 40, 1467–1477 (2022).

16. Ramirez Flores, R., Lanzer, J., Dimitrov, D., Velten, B. & Saez-Rodriguez, J. Multicellular factor analysis of single-cell data for a tissue-centric understanding of disease. eLife 12, e93161 (2023).

17. Mitchel, J. et al. Coordinated, multicellular patterns of transcriptional variation that stratify patient cohorts are revealed by tensor decomposition. Nat. Biotechnol. 43, 1192–1201 (2024).

18. Petri, M. et al. Association between changes in gene signatures expression and disease activity among patients with systemic lupus erythematosus. BMC Med. Genomics 12, 4 (2019).

19. Bombardier, C., Gladman, D. D., Urowitz, M. B., Caron, D. & Chang, C. H. Derivation of the SLEDAI. A disease activity index for lupus patients. The Committee on Prognosis Studies in SLE. Arthritis Rheum. 35, 630–640 (1992).

20. Schubert, M. et al. Perturbation-response genes reveal signaling footprints in cancer gene expression. Nat. Commun. 9, 20 (2018).

21. Müller-Dott, S. et al. Expanding the coverage of regulons from high-confidence prior knowledge for accurate estimation of transcription factor activities. Nucleic Acids Res. 51, 10934–10949 (2023).

22. Caielli, S. et al. Type I IFN drives unconventional IL-1β secretion in lupus monocytes. Immunity 57, 2497–2513.e12 (2024).

23. Urbano, P. C. M. et al. TNF-α–induced protein 3 (TNFAIP3)/A20 acts as a master switch in TNF-α blockade–driven IL-17A expression. J. Allergy Clin. Immunol. 142, 517–529 (2018).

24. Bagyinszky, E. & An, S. S. A. Genetic Mutations Associated With TNFAIP3 (A20) Haploinsufficiency and Their Impact on Inflammatory Diseases. Int. J. Mol. Sci. 25, 8275 (2024).

25. Cibrián, D. & Sánchez-Madrid, F. CD69: from activation marker to metabolic gatekeeper. Eur. J. Immunol. 47, 946–953 (2017).

26. Clark, C. E., Hasan, M. & Bousso, P. A role for the immediate early gene product c-fos in imprinting T cells with short-term memory for signal summation. PloS One 6, e18916 (2011).

27. Jonsson, A. H. et al. Granzyme K+ CD8 T cells form a core population in inflamed human tissue. Sci. Transl. Med. 14, eabo0686 (2022).

28. Guo, C.-L., Wang, C.-S., Wang, X.-H., Yu, D. & Liu, Z. GZMK+CD8+ T cells: multifaceted roles beyond cytotoxicity. Trends Immunol. 46, 562–572 (2025).

29. Blanco, P. et al. Increase in activated CD8+ T lymphocytes expressing perforin and granzyme B correlates with disease activity in patients with systemic lupus erythematosus. Arthritis Rheum. 52, 201–211 (2005).

30. Kim, J.-S. et al. IL-7Rαlow memory CD8+ T cells are significantly elevated in patients with systemic lupus erythematosus. Rheumatol. Oxf. Engl. 51, 1587–1594 (2012).

31. Xiong, H. et al. Cytotoxic CD161−CD8+ TEMRA cells contribute to the pathogenesis of systemic lupus erythematosus. eBioMedicine 90, (2023).

32. Miller, S. et al. Hypomethylation of STAT1 and HLA-DRB1 is associated with type-I interferon-dependent HLA-DRB1 expression in lupus CD8+ T cells. Ann. Rheum. Dis. 78, 519–528 (2019).

33. Viallard, J. F. et al. HLA-DR expression on lymphocyte subsets as a marker of disease activity in patients with systemic lupus erythematosus. Clin. Exp. Immunol. 125, 485–491 (2001).

34. Berard, M. & Tough, D. F. Qualitative differences between naïve and memory T cells. Immunology 106, 127–138 (2002).

35. Liberzon, A. et al. The Molecular Signatures Database (MSigDB) hallmark gene set collection. Cell Syst. 1, 417–425 (2015).

36. Psarras, A. et al. Functionally impaired plasmacytoid dendritic cells and non-haematopoietic sources of type I interferon characterize human autoimmunity. Nat. Commun. 11, 6149 (2020).

37. Gilliet, M., Cao, W. & Liu, Y.-J. Plasmacytoid dendritic cells: sensing nucleic acids in viral infection and autoimmune diseases. Nat. Rev. Immunol. 8, 594–606 (2008).

38. Crouse, J., Kalinke, U. & Oxenius, A. Regulation of antiviral T cell responses by type I interferons. Nat. Rev. Immunol. 15, 231–242 (2015).

39. Sun, J. C. & Lanier, L. L. NK cell development, homeostasis and function: parallels with CD8+ T cells. Nat. Rev. Immunol. 11, 645–657 (2011).

40. Dimitrov, D. et al. LIANA+ provides an all-in-one framework for cell-cell communication inference. Nat. Cell Biol. 26, 1613–1622 (2024).

41. Crowley, G., Alon, U. & Quake, S. R. Pareto optimality reveals an atlas of cellular archetypes. Proc. Natl. Acad. Sci. 123, e2530194123 (2026).

42. Adler, M., Korem Kohanim, Y., Tendler, A., Mayo, A. & Alon, U. Continuum of Gene-Expression Profiles Provides Spatial Division of Labor within a Differentiated Cell Type. Cell Syst. 8, 43–52.e5 (2019).

43. Hart, Y. et al. Inferring biological tasks using Pareto analysis of high-dimensional data. Nat. Methods 12, 233–235 (2015).

44. Oesinghaus, L. et al. A single-cell cytokine dictionary of human peripheral blood. 2025.12.12.693897 Preprint at 10.64898/2025.12.12.693897 (2025).

45. Alvez, M.B.et al. A human pan-disease blood atlas of the circulating proteome. Science. 390.6779 (2025)

46. Munroe, M. E. et al. Pro-Inflammatory Adaptive Cytokines and Shed Tumor Necrosis Factor Receptors are Elevated Preceding Systemic Lupus Erythematosus Disease Flare. Arthritis Rheumatol. Hoboken NJ 66, 1888–1899 (2014).

47. Mathian, A. et al. Ultrasensitive serum interferon-α quantification during SLE remission identifies patients at risk for relapse. Ann. Rheum. Dis. 78, 1669–1676 (2019).

48. Lake, B. B. et al. An atlas of healthy and injured cell states and niches in the human kidney. Preprint at 10.1101/2021.07.28.454201 (2021).

49. Chernova, I. et al. The ion transporter Na+-K+-ATPase enables pathological B cell survival in the kidney microenvironment of lupus nephritis. Sci. Adv. 9, eadf8156 (2023).

50. Sun, C.-Y. et al. The Characteristics and Significance of Locally Infiltrating B Cells in Lupus Nephritis and Their Association with Local BAFF Expression. Int. J. Rheumatol. 2013, 954292 (2013).

51. Tanevski, J., Flores, R., Gabor, A., Schapiro, D. & Saez-Rodriguez, J. Explainable multiview framework for dissecting spatial relationships from highly multiplexed data. Genome Biol. 23, 97 (2022).

52. Armingol, E., Officer, A., Harismendy, O. & Lewis, N. E. Deciphering cell–cell interactions and communication from gene expression. Nat. Rev. Genet. 22, 71–88 (2021).

53. Schäfer, P. S. L. et al. ParTIpy: A Scalable Framework for Archetypal Analysis and Pareto Task Inference. 2025.09.08.674797 Preprint at 10.1101/2025.09.08.674797 (2025).

54. Barturen, G., Beretta, L., Cervera, R., Van Vollenhoven, R. & Alarcón-Riquelme, M. E. Moving towards a molecular taxonomy of autoimmune rheumatic diseases. Nat. Rev. Rheumatol. 14, 75–93 (2018).

55. Samões, B., Zen, M., Abelha-Aleixo, J., Gatto, M. & Doria, A. Caveats and pitfalls in defining low disease activity in systemic lupus erythematosus. Autoimmun. Rev. 21, 103165 (2022).

56. Altabás-González, I. et al. Validation of proposals for definitions of moderate and severe disease activity in SLE: impact on flares, quality of life, damage accrual, hospitalisations and mortality. Lupus Sci. Med. 12, e001766 (2025).

57. Fanouriakis, A. et al. EULAR recommendations for the management of systemic lupus erythematosus with kidney involvement: 2025 update. Ann. Rheum. Dis. 85, 75–90 (2026).

58. Fanouriakis, A. et al. EULAR recommendations for the management of systemic lupus erythematosus: 2023 update. Ann. Rheum. Dis. 83, 15–29 (2024).

59. Baker, T. et al. Type I interferon blockade with anifrolumab in patients with systemic lupus erythematosus modulates key immunopathological pathways in a gene expression and proteomic analysis of two phase 3 trials. Ann. Rheum. Dis. 83, 1018–1027 (2024).

60. Ramsköld, D. et al. B cell alterations during BAFF inhibition with belimumab in SLE. eBioMedicine 40, 517–527 (2019).

61. Scherlinger, M., Kolios, A. G. A., Kyttaris, V. C. & Tsokos, G. C. Advances in the treatment of systemic lupus erythematosus. Nat. Rev. Drug Discov. 24, 926–944 (2025).

62. Wolf, F., Angerer, P. & Theis, F. SCANPY: large-scale single-cell gene expression data analysis. Genome Biol. 19, 15 (2018).

63. Robinson, M. D., McCarthy, D. J. & Smyth, G. K. edgeR: a Bioconductor package for differential expression analysis of digital gene expression data. Bioinformatics 26, 139–140 (2010).

64. Eadon, M. T. et al. Kidney Histopathology and Prediction of Kidney Failure: A Retrospective Cohort Study. Am. J. Kidney Dis. Off. J. Natl. Kidney Found. 76, 350–360 (2020).

65. Stuart, T. et al. Comprehensive Integration of Single-Cell Data. Cell 177, 1888–1902.e21 (2019).

66. Martín-Fernández, J. A., Barceló-Vidal, C. & Pawlowsky-Glahn, V. Dealing with Zeros and Missing Values in Compositional Data Sets Using Nonparametric Imputation. Math. Geol. 35, 253–278 (2003).

67. Muzellec, B., Teleńczuk, M., Cabeli, V. & Andreux, M. PyDESeq2: a python package for bulk RNA-seq differential expression analysis. Bioinformatics 39, btad547 (2023).

68. Badia-I-Mompel, P., et al. decoupleR: ensemble of computational methods to infer biological activities from omics data. Bioinforma. Adv. 2, vbac016 (2022).

69. Argelaguet, R. et al. MOFA+: a statistical framework for comprehensive integration of multi-modal single-cell data. Genome Biol. 21, 111 (2020).

70. Dimitrov, D. et al. Comparison of methods and resources for cell-cell communication inference from single-cell RNA-Seq data. Nat. Commun. 13, 3224 (2022).

71. Therneau, T. M. A Package for Survival Analysis in R. R package version 3.5–8 (2024). https://CRAN.R-project.org/package=survival

